# The recent evolutionary rescue of a staple crop depended on over half a century of global germplasm exchange

**DOI:** 10.1101/2021.05.11.443651

**Authors:** Kebede T. Muleta, Terry Felderhoff, Noah Winans, Rachel Walstead, Jean Rigaud Charles, J. Scott Armstrong, Sujan Mamidi, Chris Plott, John P. Vogel, Peggy G. Lemaux, Todd C. Mockler, Jane Grimwood, Jeremy Schmutz, Gael Pressoir, Geoffrey P. Morris

## Abstract

Rapid environmental change can lead to extinction of populations or evolutionary rescue via genetic adaptation. In the past several years, smallholder and commercial cultivation of sorghum (*Sorghum bicolor*), a global cereal and forage crop, has been threatened by a global outbreak of an aggressive new biotype of sugarcane aphid (SCA; *Melanaphis sacchari*). Here we characterized genomic signatures of adaptation in a Haitian sorghum breeding population, which had been recently founded from admixed global germplasm, extensively intercrossed, and subjected to intense selection under SCA infestation. We conducted evolutionary population genomics analyses of 296 post-selection Haitian lines compared to 767 global accessions at 159,683 single nucleotide polymorphisms. Despite intense selection, the Haitian population retains high nucleotide diversity through much of the genome due to diverse founders and an intercrossing strategy. A genome-wide fixation (*F*_ST_) scan and geographic analyses suggests that adaptation to SCA in Haiti is conferred by a globally-rare East African allele of *RMES1*, which has also spread to other breeding programs in Africa, Asia, and the Americas. *De novo* genome sequencing data for SCA resistant and susceptible lines revealed putative causative variants at *RMES1*. Convenient low-cost markers were developed from the *RMES1* selective sweep and successfully predicted resistance in independent U.S. × African breeding lines and eight U.S. commercial and public breeding programs, demonstrating the global relevance of the findings. Together, the findings highlight the potential of evolutionary genomics to develop adaptive trait breeding technology and the value of global germplasm exchange to facilitate evolutionary rescue.

## INTRODUCTION

Ongoing processes of global change, encompassing climate change, nutrient cycles, and pest outbreaks, are shaping the evolution of natural and agricultural ecosystems (*1, 2*). Intense selection pressure following environment changes may lead to the rapid decline or extinction of populations (*3, 4*). If a population is to persist under such strong selection, adaptive standing genetic variation must exist or adaptive *de novo* variation must arise on a sufficiently fast timescale (*5*). This population genetic phenomenon, evolutionary rescue, has become a focus of considerable empirical and theoretical study in ecology and conservation biology, since the current rate of global change could exceed the capacities of many populations to adapt (*6, 7*). Still, there is a lack of examples of evolutionary rescue occuring in the field and at large geographic scales (*4*). In agricultural systems, the spread of pests or emergence of new aggressive biotypes may lead to a reduction of crop diversity or a total loss of crop cultivation (*8*). Therefore, understanding and facilitating evolutionary rescue in agricultural systems is critical for global food security.

Populations of crops or wild species subjected to strong selection pressure may experience a major population bottleneck, resulting in a loss of genetic diversity (*9*). The level of diversity preserved in a population recovering from strong selection depends on the number of backgrounds on which the adaptive alleles emerge (*10*), which can determine the potential for future adaptation or genetic gain. Conversely, adaptation conferred by a beneficial variant derived from a single progenitor causes the removal of genetic diversity from the surviving population (*10, 11*). Evolutionary population genomics approaches using genome-wide polymorphism data from diverse germplasm can identify candidate loci for adaptive traits (*12*). While genome scans for selection have been widely used to identify putative adaptive alleles in crops (*9, 13*), they have not yet been used to identify trait-predictive markers for molecular breeding of stress-resilient varieties (*14*).

Sorghum (*Sorghum bicolor* L. [Moench]) is among the world’s most important staple crops for smallholder farmers in semiarid regions, as well as a commercial grain and forage crop in industrialized nations (*15*). Since 2013 an aggressive biotype of the sugarcane aphid (SCA; *Melanaphis sacchari*) has become a major threat to global sorghum production, with widespread and substantial yield loss (*16, 17*). The *M. sacchari* superclone has been rapidly expanding (*18*), putting >90% of the sorghum-producing areas of North America at risk and threatening to end sorghum cultivation in some areas (*16*). In Haiti, a Caribbean nation with one of the world’s highest rates of food insecurity, sorghum is among the most important staple crops (*19*). However, heavy infestations by *M. sacchari* since 2015 have caused the loss of over 70% of sorghum production in the country and prevented production of most local landraces (*20*). Shortly before the SCA outbreak, a new Haitian breeding population had been launched by Chibas using global admixed germplasm, rapid-cycling intercrossing, and selection under smallholder conditions (i.e. no insecticidal treatment) (*21*). Selecting from a small number of breeding lines that survived SCA infestation, a new SCA resistant sorghum variety, Papèpichon, was developed and distributed nationally (*19*), and intercrossing and advancement of resistant breeding lines has continued.

Here we used a retrospective genomic analysis of the Haitian sorghum breeding population that was subjected to strong selection under SCA infestation, to understand the genetic basis of the evolutionary rescue following the SCA outbreak, as well as the origins of the SCA resistance alleles. We find that the rapid adaptation of the Haitian breeding population to the SCA outbreak was due to selection for a globally-rare Ethiopian allele at the *RMES1* SCA resistance locus, which is shared across programs in Africa, Asia, and the Americas because of >50 years of global germplasm exchange prior to the SCA outbreak. Further, we developed a convenient low-cost molecular marker based on the evolutionary genome scan and validated it in eight commercial and public sorghum breeding programs, demonstrating the value of leveraging global germplasm exchange and evolutionary population genomics to improve crop resilience.

## RESULTS

### Genome-wide polymorphism and nucleotide diversity

To understand the evolutionary rescue of sorghum following the SCA outbreak, we conducted a retrospective genomic analysis of the Haitian breeding population (HBP) in comparison to a global diversity panel (GDP). Genotyping-by-sequencing of 296 HBP and 767 GDP (Supp. Fig. S1; Supp. File S1) sorghum lines generated 159,683 polymorphic SNPs with an average SNP density of 75 and 229 per Mb in the HBP and GDP, respectively (Supp. Fig. S2). The GDP had a higher proportion of low-frequency minor alleles (<5% MAF) compared to the HBP (Supp. Fig. S3). Average inbreeding coefficients (*F*_IS_) in HBP and the GDP was estimated at 0.7 and 0.9, respectively (Supp. Table S1). The effect of selection on genetic diversity in HBP was assessed based on genome-wide nucleotide diversity (π) in the HBP in comparison to (i) the GDP and (ii) a major public program in the US (Texas A&M pre-breeding lines, TAM-PBL, *N* = 35). Average nucleotide diversity in the HBP was estimated at 2.3×10^−5^. In the GDP and TAM-PBL, estimates of average π were 5.8×10^−5^ and 4.8×10^−5^, respectively (Fig. 1A-C, Supp. Table S2). In the HBP, 31% of 1 Mb windows have negative average Tajima’s *D* values, while in the GDP predominantly positive values of Tajima’s *D* were observed (Supp. Fig. S4).

**Figure 1:**
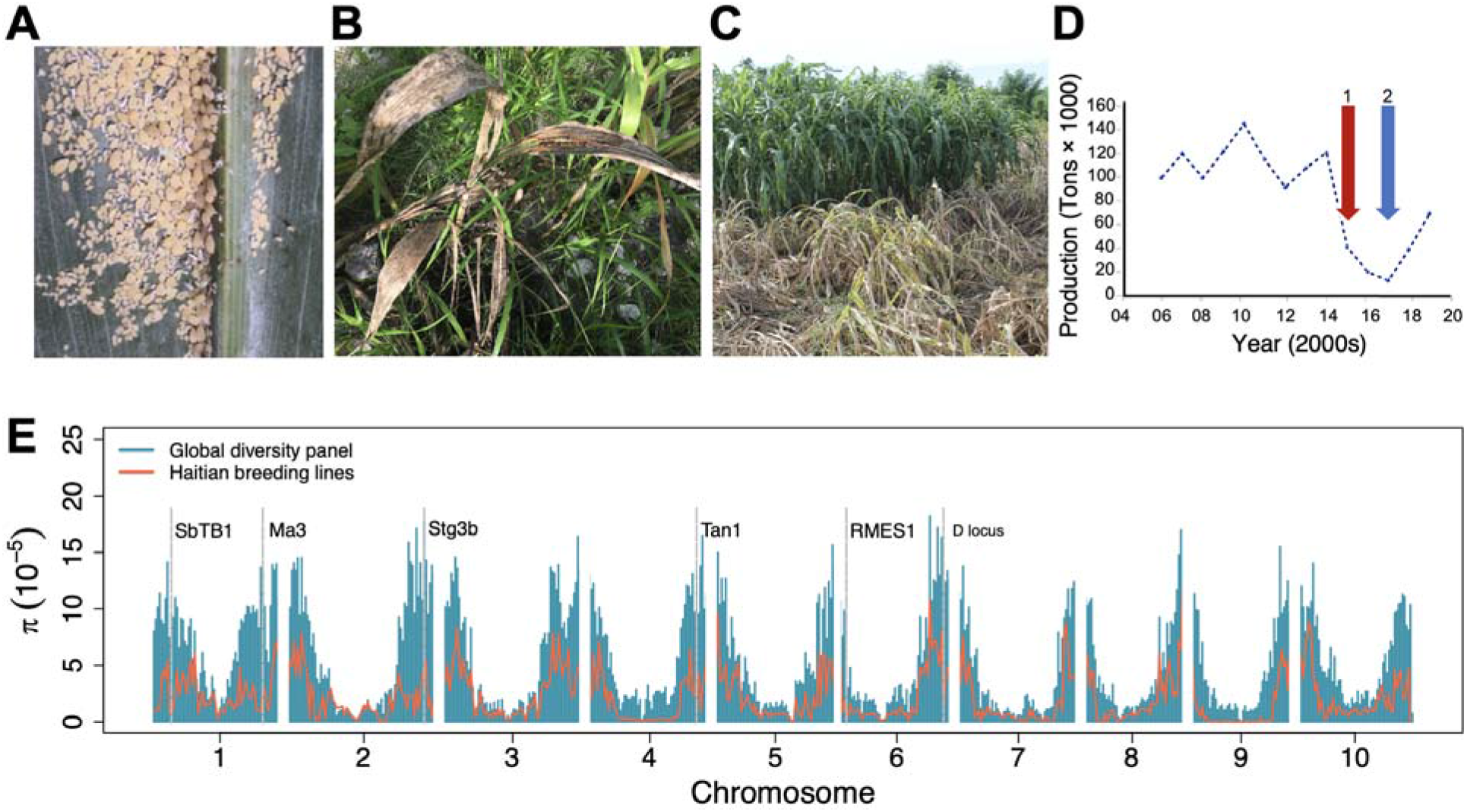
Evolutionary rescue following a continental outbreak of a sorghum pest. (A) Infestation of sugarcane aphid (SCA), *M. sacchari*, on a commercial hybrid in the US sorghum-growing production region (Kansas). (B) SCA infestation on a traditional sorghum variety on a smallholder farm in Haiti (brown plant in foreground; green leaves in background are maize and wild grasses). (C) Reaction of susceptible (brown plants; foreground) and resistant (green plants; background) sorghum breeding lines under natural SCA infestation during breeding trials in Haiti. (D) Estimates of annual sorghum production in Haiti (2006-2019), indicating the start of the SCA outbreak (1, red arrow) and the start of national distribution of SCA resistant variety, Papépichon (2, blue arrow). (E) Genome-wide nucleotide diversity (π) in the Haitian breeding population (red line) compared to a global diversity panel (blue bars). Nucleotide diversity was calculated for a non-overlapping sliding window of 1 Mbp across the genome. The grey vertical dashed lines indicate the position of *a priori* candidate genes for breeding targets of the Haiti program which colocalized with genomic regions of reduced π (see Supp. File S3 for details).

### Contributions of global sorghum diversity to the Haitian breeding population

The genetic ancestry of the HBP from global germplasm was inferred based on population structure analyses. In a neighbor joining analysis, the HBP clusters with caudatum accessions (Fig. 2A), specifically caudatums from East Africa. Similarly, in principal coordinate analysis, the HBP cluster swith East African caudatum accessions (Fig. 2C). To estimate ancestry coefficients for HBP lines, we used Bayesian model-based clustering in ADMIXTURE, projecting HBP lines onto ancestral populations and allele frequencies defined using only GDP (with HBP lines omitted). With the GDP, the lowest cross-validation error was observed at *K* = 8 (Supp. Fig. S5) and accessions clustered by ecogeographic region and botanical type, as expected. ADMIXTURE projection analysis suggests that the HBP is admixed, largely consisting of caudatum haplotypes (>80% of the genome) with a remaining small percentage being contributed by durra and guinea sorghums (Fig. 2D).

**Figure 2:**
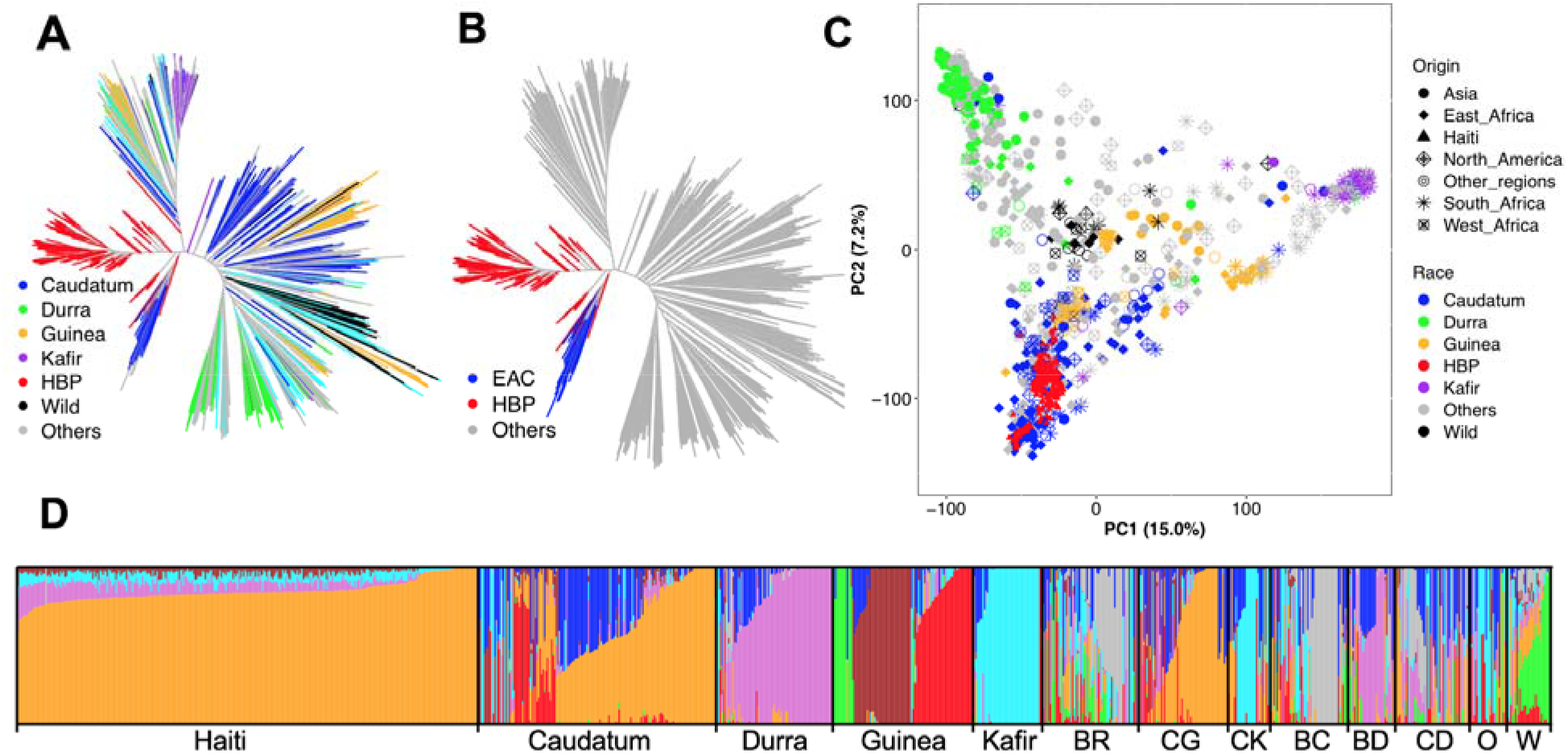
Population structure of the Haitian breeding population in relation to global sorghum diversity reflects its derivation from East African germplasm. Genetic relatedness of the Haitian breeding population (HBP) to the global diversity assessed by neighboring joining method, color-coded by botanical type (A) or highlighting the close relationship between the HBP and East African caudatum (EAC) germplasm (C) Scatterplot of the first two principal component (PC) of genome wide SNP variation, demonstrating the clustering of HBP within EAC germplasm. (D) Bayesian hierarchical clustering of the HBP and GDP with the probability of membership (*Q*) in each of K = 8 ancestral populations. The Q-value bar plots are arranged by botanical types to reflect the relationship of the HBP to the GDP. Note, color-coding of the bar plots in panel D is arbitrary and does not reflect the color-code in panels A-C. BR = Bicolor, CG = caudatum-guinea, CK= caudatum-kafir, BC = bicolor-caudatum, BD = bicolor-durra, CD = caudatum durra, O = others (includes botanical types containing less than 10 individuals), W= wild.

### Evidence of a selective sweep in the Haitian breeding population at *RMES1*

To identify genome regions implicated in the evolutionary rescue of the HBP, genome-wide scans for outlier loci were performed based on an *F*_ST_ test. Overall, the HBP is moderately differentiated from the global diversity panel, with an average genome-wide *F*_ST_ of 0.16 (Fig. 3A, Supp. File S2). Based on a Bonferroni-adjusted *P*-value < 0.01, *F*_ST_ analysis identified 171 outlier genomic regions, which are candidate selective sweep regions. Several genomic regions with *F*_ST_ outlier regions co-localized with candidate genes for traits under selection by the Chibas breeding program, including photoperiodic flowering, inflorescence architecture, stay□green, stem sugar content, and SCA resistance (Fig 1D; Supp. File S3). Interestingly, the most extreme *F*_ST_ outliers were observed on chromosome 6, precisely colocalizing with *RMES1*, a locus previously shown to underlie SCA resistance in a Chinese sorghum line of unknown pedigree (*22*) (Fig. 3A-B). To characterize the prevalence of the putative selected haplotype and identify its geographic origin, we mapped the allelic distribution of the highest *F*_ST_ SNP S6_2995581 in global georeferenced sorghum landraces (Fig. 3C) and compared these distributions to the allele frequency in US and Haitian breeding germplasm (Fig. 3C, inset left). Globally, the allele is rare (<2%), found only in Ethiopian caudatum landraces and a few breeding lines from West Africa and the US. However, the sweep-associated allele is common (∼40%) in Ethiopian caudatum accessions (Fig. 3C; Supp. Table S3). The high local frequency of the sweep-associated allele in Ethiopia suggests a likely origin of the SCA resistance allele in the Ethiopian highlands (Fig. 3C, inset right).

**Figure 3:**
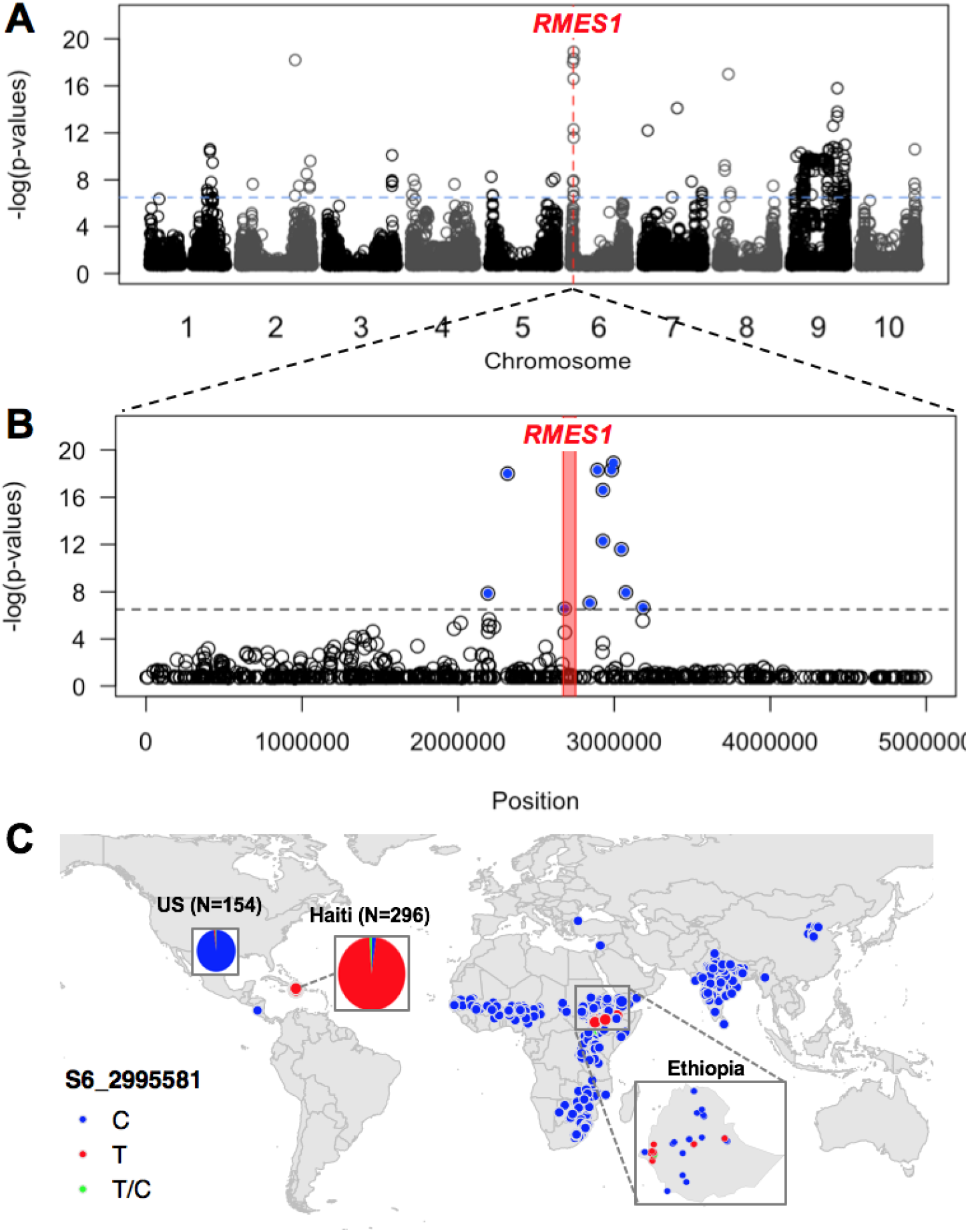
Genome scan for selection identifies the major aphid resistance allele at *RMES1* originating in Ethiopia. (A) Genome-wide scan for selection in the Haitian breeding population using fixation index (*F*_ST_) with the -log(*F*_ST_ *p*-value) (y-axis) plotted against position on the chromosome (x-axis). (B) Detailed view (5 Mb) of top *F*_ST_ peak on chromosome 6 that colocalizes with the *RMES1* locus. The ∼130 kb region from 2,667,082 to 2,796,847 bp corresponding to the published *RMES1* interval is denoted with the red bar. (C) Global allele distribution of the SNP that showed the highest *F*_ST_ value (S6_2995581), which colocalized with the *RMES1* locus. Allelic state for georeferenced global germplasm is denoted with points. Allele frequencies in the United States (C=151, T=2, T/C=1) and Haiti (C=6, T=287, T/C =3) breeding germplasm, denoted in pie charts with area proportional to number of accessions, show the allele is almost fixed in Haitian breeding germplasm and rare in U.S. breeding germplasm.

### Comparative genomic analysis to identify candidate causative variants

To identify candidate causative variants for the *RMES1* locus, we used whole-genome resequencing and *de novo* genome sequencing of sorghum accessions with known SCA reactions. The *RMES1* interval previously defined based on biparental linkage mapping (*22*) includes seven gene models (Sobic.006G017000, Sobic.006G017100, Sobic.006G017200, Sobic.006G017332, Sobic.006G017266, Sobic.006G017400, and Sobic.006G017500) that were candidates for the causative gene. Comparative genomic analyses based on local multiple sequence alignment (MSA) of *de novo* genome sequence of the resistant accession (PI 276837, the Ethiopian progenitor of SCA resistant line SC170) and three sorghum reference genomes of SCA susceptible lines (BTx623, Tx430, and BTx642) were used to identify potential causative variants. No sequence variants were identified in the exons of three of the seven genes (Sobic.006G017000, Sobic.006G017100, and Sobic.006G017266). A total of 35, 32, and 29 nonsynonymous SNPs were detected in the exons of Sobic.006G017200, Sobic.006G017400, and Sobic.006G017500, when comparing the sequences of the resistant PI 276837 and the three susceptible accessions. In addition, three insertion-deletion variations resulting in frame-shift were detected in Sobic.006G017500. (Supp. File S4). To further refine the set of candidate causative variants, we performed a localized association analysis for SCA resistance (“resistant” or “susceptible”, based on literature classification) around *RMES1* with resequencing data for diverse sorghum accessions (Fig. 4, Supp. File S5) that detected 101 highly significant associations (*P*-value > 0.0001). Annotations of the variants within the *RMES1* locus indicate that only ten of 101 associated variants are nonsynonymous (5 of 10 in Sobic.006G017200 and the remaining 5 of 10 in Sobic.006G017500.

**Figure 4:**
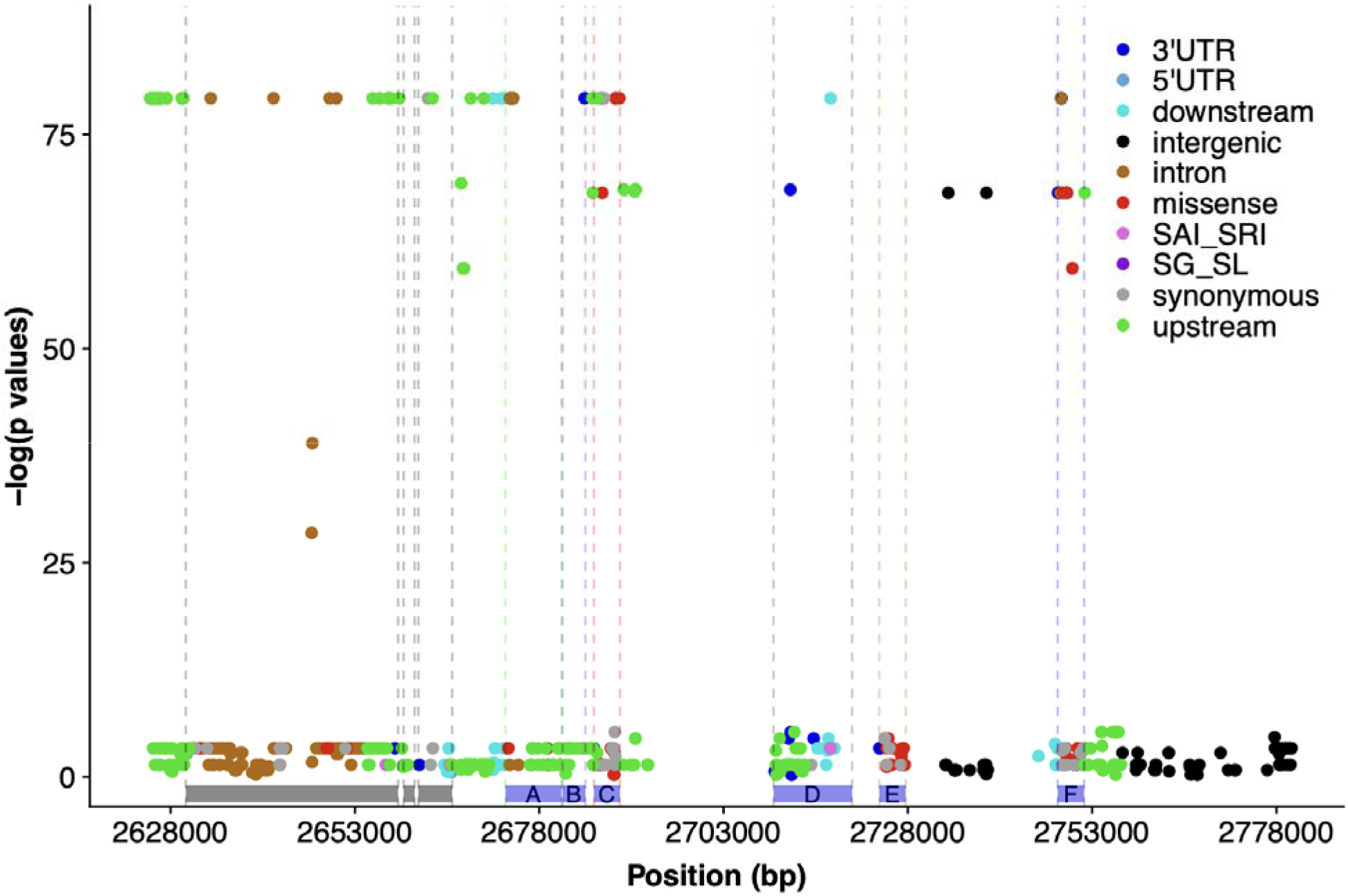
Whole-genome resequencing and local association mapping identifies potential causative variants at *RMES1*. Functional annotation and association mapping of nucleotide polymorphisms within the *RMES1* locus across a set of 13 diverse sorghum accessions with known SCA resistance or susceptibility. The -log of *p*-values of local marker-trait association scan plotted against the chromosomal positions at the *RMES1* locus on chromosome 6. Variants are color-coded by annotation generated by the SnpEff program. Blue bars represent the seven annotated genes within the *RMES1* interval (A = Sobic.006G017000, B = Sobic.006G017100, C = Sobic.006G017200, D = Sobic.006G017332 and Sobic.006G017266, E = Sobic.006G017400, F = Sobic.006G017500.v3.1). Grey bars indicate genes outside of the *RMES1* interval as originally defined (*22*). 3’UTR: 3 prime UTR variant, 5’UTR: 5 prime UTR variant, downstream: Downstream gene variant, intergenic: intergenic region, intron: Intron variant, missense: Missense variant, SAI_SRI: splice acceptor/intron or splice region intron variants, SG_SL: stop gained or stop loss variant, synonymous: Synonymous variant upstream: Upstream gene variant

### Development and validation of a molecular marker based on the selective sweep

Next, we sought to test the hypothesis that the genome region identified by the *F*_ST_ scan underlies variation for SCA resistance in other global sorghum germplasm. Therefore we developed a kompetitive allele specific PCR (KASP) marker based on the SNPs at the *RMES1* locus identified in the *F*_ST_ scan. Of the candidate SNPs (Supp. File S2, Supp. Table S3, Supp. File S6), SNP 06_02892438 was determined to have the best combination of linkage, LD, and technical KASP functionality of the SNPs. Alternative SNPs were also developed into markers (Supp. File S6), and while the markers are often used as technical checks, testing has confirmed the priority of the marker based on SNP 06_02892438 (Sbv3.1_06_02892438R). Initial validation of the Sbv3.1_06_02892438R KASP marker using DNA samples from known resistant lines (SC110, Tx2783, and IRAT204), susceptible lines (BTx623 and BTx642) (*23*), and multiple F_2_ families segregating for SCA resistance demonstrated that the KASP marker Sbv3.1_06_02892438R was in complete agreement with historical phenotypes of inbred lines and segregated within F_2_ populations (Supp. File S7).

An F_4_ population derived from a cross between IRAT204 (resistant African variety) and Tx430 (susceptible US breeding line) was used to further validate the broader utility and predictiveness of the KASP marker for marker-assisted selection (Fig. 5A-B). A total of 50 F_4_ lines together with resistant (IRAT204 and SC110) and susceptible controls (RTx430) were genotyped with the KASP marker Sbv3.1_06_02892438R. Both resistant controls and 23 F_4_ lines were homozygous for the resistant allele. The susceptible control and 9 F_4_ lines were homozygous for the susceptible allele, and the remaining 18 F_4_ lines were heterozygous at the SNP. Twenty-three selected F_4_ lines with three resistant and three susceptible control lines were tested for SCA reaction in a free-choice flat screen assay in the greenhouse, scoring aphid damage rating, leaf greenness (SPAD), and seedling height. The SCA reaction phenotypes match the KASP marker genotypes, demonstrating the reliability and predictability of using KASP markers in marker-assisted selection for SCA resistance breeding (Fig. 5A-B; Supp. File S7).

**Figure 5:**
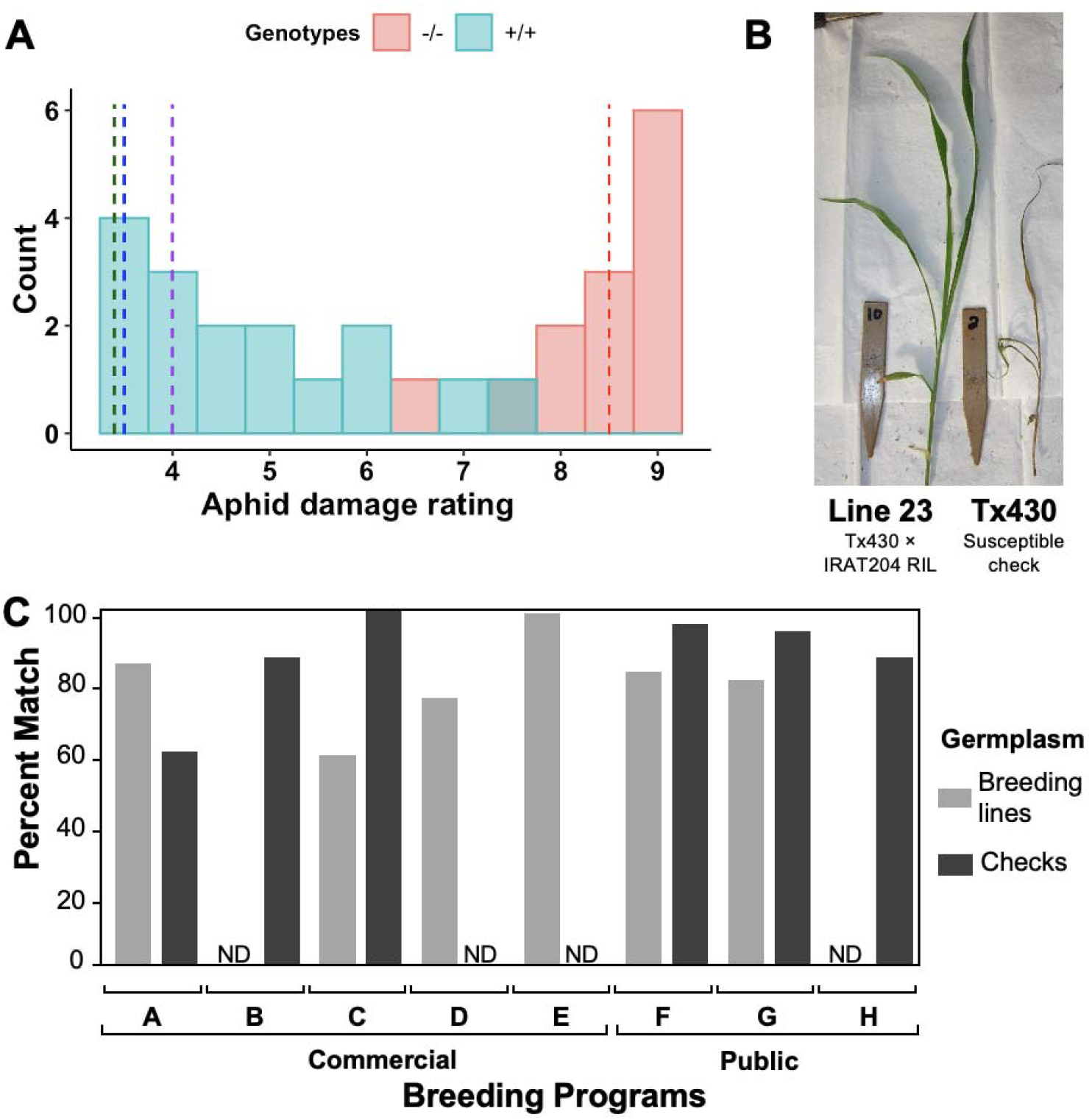
Multi-program evaluation of a molecular marker developed based on the selective sweep validates its global utility. (A) The KASP marker predicts SCA resistance in independent US × Senegal breeding lines. The histogram represents the aphid damage ratings of F_4_ lines from a Tx430 × IRAT204 (*N* = 22) family under infestation by *M. sacchari* at the seedling stage in a choice greenhouse assay. The cyan bars represent the aphid damage ratings for lines carrying the +/+ genotypes at the SNP 06_02892438, while the red bars represent aphid damage ratings of the lines carrying the -/- genotypes. The blue dashed lines represent the average aphid damage rating of the resistant checks Tx2783, IRAT204, and SC110 (green, blue, and purple dashed lines, respectively) while red dashed line represents the average damage rating of both susceptible checks, Tx7000 and Tx430. (B) Representative SCA reaction from the choice greenhouse assay for an F4 line carrying the +/+ genotype (left) versus the susceptible parent Tx430 (right). (C) Evaluation of the same marker in eight US breeding programs. Percent match of KASP marker genotyping prediction with breeder-provided SCA resistance classification for five commercial breeding programs and three public breeding programs. ND = Not determined.

### Multi-program validation and deployment in commercial and public breeding programs

To further validate the utility of the SCA resistance KASP markers, we tested them with five US commercial seed company breeding programs and three US public sector breeding programs, representing a large fraction of the US sorghum breeding community (Fig. 5C). (The programs are anonymized to avoid disclosing proprietary information.) Under the hypothesis that (i) *RMES1* underlies SCA resistance in US breeding programs and (ii) the KASP marker (Sbv3.1_06_02892438R) tags the relevant resistant vs. susceptible haplotypes, the breeders’ phenotype-based classification of SCA resistance should largely match the KASP marker genotype-based prediction. As expected, the match between the phenotype-based breeder classification and KASP marker genotypes is high, ranging from ∼60-100%, with most germplasm sets (9/12) have >80% matching (Fig. 5C; Fig. S6). Less than 0.5% of mismatches (5/1100) were observed among technical replicates (independent tissue samples from the same plant), so mismatches are unlikely to be due to KASP genotyping errors. Mismatches may be due to differences among programs of SCA resistant or susceptible haplotypes, or errors in the phenotype-based resistance classifications (some of which are based on visual ratings under natural field infestations, which are prone to false positives (*24*)). There were also some genotype-phenotype mismatches in public germplasm checks used by commercial and public programs (Fig. 5C). In nearly all cases, further investigation revealed that mismatches were due to unexpected heterogeneity in public germplasm within or among breeding programs (Supp.Table S4).

## DISCUSSION

### *RMES1* is a major resistance gene underlying evolutionary rescue of sorghum worldwide

Understanding the genetics of evolutionary rescue, including the genetic architecture and molecular basis, could contribute to more resilient conservation and breeding strategies (*25*). Here we hypothesized, parsimoniously, that a single Mendelian SCA resistance locus *RMES1* could underlie the global evolutionary rescue of sorghum to the new *M. sacchari* superclone. Previous studies had suggested that a single dominant locus is responsible for SCA resistance in families derived from resistant Chinese grain sorghum variety Henong 16 (H16) and susceptible BTx623, or families derived from US breeding lines, resistant RTx2738 and susceptible CK60 (*22, 26*). The H16 resistance was mapped to a ∼130 kb region at 2.7 Mb on chromosome 6 (*RMES1*) (*22*). Consistent with the *RMES1* evolutionary rescue hypothesis, the genome region with the highest *F*_ST_ in the HBP colocalized precisely with *RMES1* (Fig. 3). Together, the evolutionary genome scan (Fig. 3) and multi-program marker validation (Fig. 5) provides strong evidence that *RMES1* is the major SCA resistance locus globally, shared across the Americas, Asia, and Africa. However, our findings do not preclude the hypothesis that other SCA resistance loci were selected in Haiti and were required for the evolutionary rescue. In particular, other *F*_ST_ scan peaks on chromosome 2, 7, 8, and 9 (Fig. 3) could correspond to other SCA resistance loci. Given that SCA resistance is fixed in the Haitian program, further population development and quantitative trait locus mapping for SCA resistance will be necessary to test this hypothesis.

Identifying the causal variant underlying SCA resistance would advance our understanding of aphid resistance mechanisms in plants (*27*) and facilitate development of perfectly-predictive molecular markers for SCA resistance breeding (*28*). Our comparative genomic analysis between the resistant PI 276837 and the three susceptible reference genomes identified four candidate genes with putative functional variants within the *RMES1* locus (Sobic.006G017200, Sobic.006G017332, Sobic.006G017400 and Sobic.006G017500; Supp. File S4). Three of the four genes in the candidate region encode leucine-rich repeat (LRR) proteins, a gene family involved in immune responses to invading pathogens and insects (*29*). Given that some LRR genes mediate plant resistance to aphids and other phloem-feeding insects (*27*) these genes represent promising candidates for the *RMES1* causative gene. Functional annotation and sequence comparison between the resistant and susceptible accession identified non-synonymous variants only in Sobic.006G017200 and Sobic.006G017500 (Fig. 4), suggesting these NLR are promising candidates for the *RMES1* gene. Fine-mapping and positional cloning will be needed to test these hypotheses and positively identify the causative variant.

### Evolutionary rescue of sorghum depended on a half century of global germplasm exchange

In the twentieth century, sorghum genebanks and breeding programs exchanged germplasm widely (*30, 31*). Based on pedigree records and morphology we hypothesized that the Haitian breeding population originated from global admixed germplasm with a primary contribution of Ethiopian caudatum of the zerazera working group. Consistent with this hypothesis, HBP genotypes clustered with caudatum sorghum of East Africa (Fig. 2), but admixture analysis identified a contribution from durra and guinea sorghum from West Africa (Fig. 2D). Combining population genomics findings (Fig. 2, 3) with genebank and pedigree records (*32, 33*), we can map the history of global germplasm exchange that led to the evolutionary rescue of sorghum in Haiti following the SCA outbreak (Fig. 6A), as well as the spread of the SCA resistance allele from Ethiopia to breeding programs around the world (Fig. 6B). Notably, the evolutionary rescue of sorghum in the Americas (Haiti and US) involved germplasm and knowledge exchange over a period of >50 years, involving nine countries on three continents.

**Figure 6:**
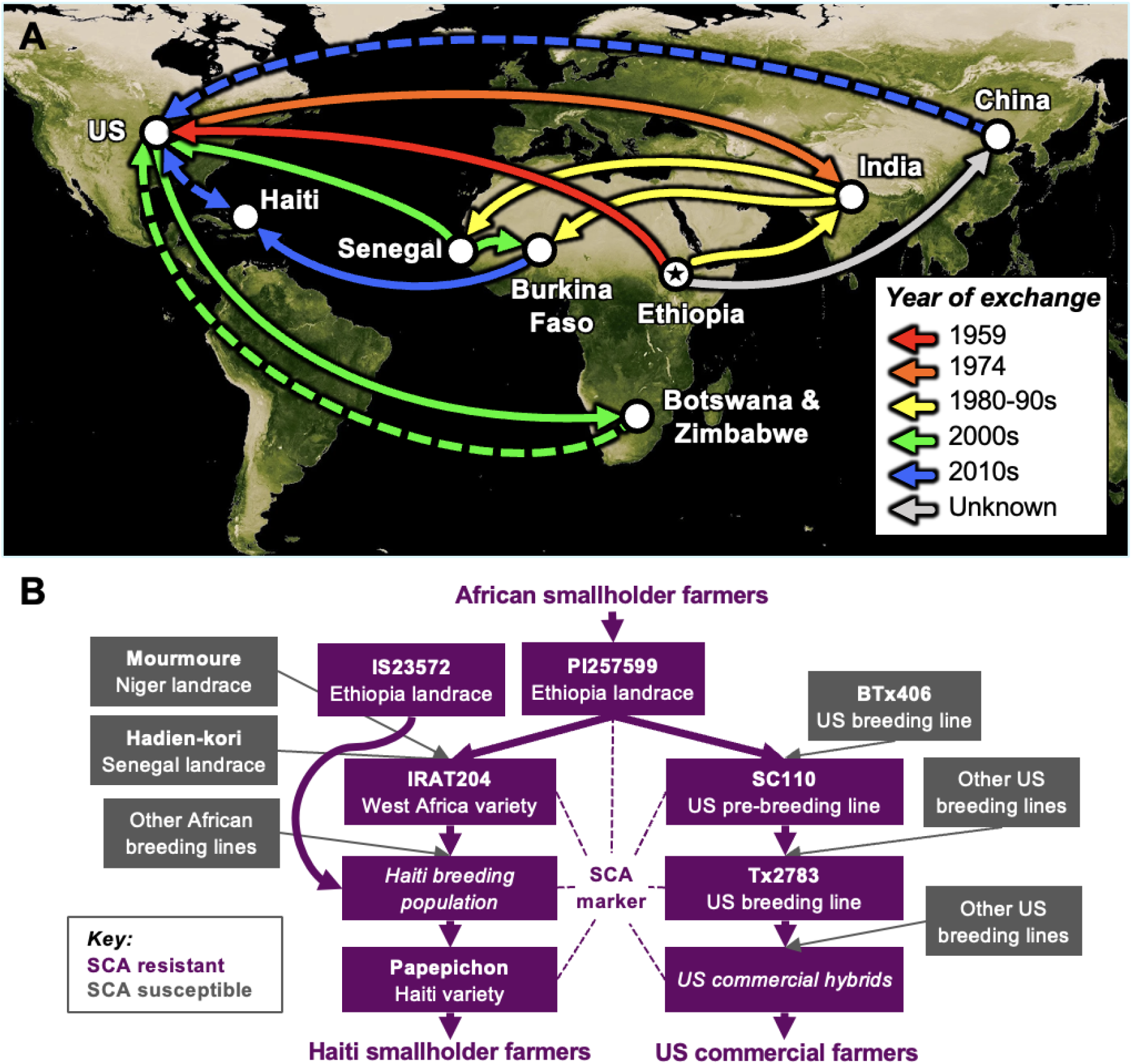
Evolutionary rescue of sorghum through >50 years of global exchange of germplasm and knowledge. (A) Germplasm and knowledge exchange inferred from pedigree records and genomic analyses. Germplasm exchange is denoted by solid lines. Knowledge exchange through scientific literature is denoted in dashed lines. The star indicates the inferred origin of the SCA resistance allele in the Ethiopian highlands, with at least two paths to the Americas, via IS 23572 (yellow line) or PI 257599 (red line). (B) Pedigree relationships among global accessions, breeding lines, breeding programs, or varieties, color-coded by inferred SCA resistance or susceptibility. Note, with respect to US commercial hybrids, the diagram is illustrative and is not meant to imply that all US commercial hybrids used Tx2783 as the SCA resistance donor. Some known pedigree information has been omitted from the diagram for clarity.

In the case of the SCA outbreak, the global sorghum improvement community was fortunate that the rare SCA resistance allele originated in East African caudatums, since this germplasm is preferred by many sorghum breeders worldwide and widely used by breeding programs in Africa, Asia, and the Americas (*30, 34*). The SCA resistance allele appears to have been inadvertently spread across sorghum breeding programs across the three continents long before the recent SCA outbreak (Fig. 6). For example, SC110, a converted version of an Ethiopia caudatum landrace (PI 257599/IS 12610) identified as SCA resistant in several world regions (*23, 35*), is a major contributor to the pedigrees of most SCA-resistant breeding lines in the US (Fig. 6B) (*36*). The same progenitor line (IS 12610) was used by breeding programs in West Africa (Fig. 6B) as a parent of IRAT204 (CE151-262; PI 656031), a widely-adopted variety (*37*) and key progenitor of current West African breeding programs (*34*).

Another potential benefit of germplasm exchange is the maintenance of diversity in breeding programs following strong selection, including evolutionary rescue. Given the strong selection on the HBP during the SCA outbreak, it might be expected that the post-selection HBP no longer retains sufficient diversity for future adaptation and genetic gain (*7*). However, the HBP was founded with diverse admixed global germplasm (Fig. 2) and extensively intercrossed, so it appears to have retained sufficient genetic diversity for future adaptation and crop improvement. We observe only a modest reduction in nucleotide diversity observed throughout the genome of the HBP relative to global accessions, East African caudatums, or a major public pre-breeding program (Fig. 1E; Supp. Fig. S7). Recombination during intercrossing cycles (prior to the SCA outbreak) presumably reshuffled the SCA resistance allele onto many backgrounds, suggesting that the intercrossing approach was critical to allow the Haitian program to retain diversity for future genetic gain and adaptation.

### Rapid discovery and deployment of a global trait-predictive molecular marker using evolutionary population genomics

Molecular marker development based on phenotype-to-genotype mapping of trait loci (e.g. linkage or association mapping) is limited by availability of suitable mapping populations, phenotyping capacity, and genotyping resources, which can take years to develop (*13, 38*). For instance, spatial and temporal variability of SCA pressure in field trials limits the effectiveness of field phenotyping (*24*), while greenhouse assays can be complicated and time-consuming for lower-resourced programs. Thus, an evolutionary genomics approach, which leverages a history of selection by smallholder farmers or plant breeders, could have advantages for marker discovery. Despite wide use of evolutionary genome scans in crops, the hypotheses generated on adaptive loci are rarely, if ever, tested by independent experimental approaches (e.g. with near isogenic lines) (*39*). To our knowledge, this is the first example where an evolutionary or population genomic scan led directly to molecular breeding technology in use in commercial and public varietal development (Fig. 3, 5).

Here we demonstrated the effectiveness of the evolutionary population genomic approach, showing that a marker discovered in a single developing-country breeding program (Chibas-Haiti) can link crop improvement efforts across three continents (North America, Africa, Asia; Fig. 5A, 6) and across the commercial and public sector (Fig. 5A, C). Thus, our findings establish the value of evolutionary population genomics to facilitate and guide global crop improvement. The KASP marker developed and validated in this study can facilitate the rapid conversion of existing farmer-preferred varieties for SCA resistance (e.g. by marker-assisted introgression) (*40*). While the *RMES1* resistance allele is currently conferring effective resistance, a further biotype shift in the aphid could overcome this gene. Several biotype shifts occurred in the 1960-1980s for the greenbug aphid *Schizaphis graminum* (*41*) and slowed genetic gain in sorghum for many years (*42*). The markers developed here could facilitate identification of new SCA resistance genes, via by counterselection of *RMES1* allele to reveal novel SCA resistance. These outsourced KASP markers are convenient for breeding programs, since they require no laboratory labor or facilities, and are low cost relative to dedicated field or greenhouse phenotyping capacity, at ∼$2 per sample for DNA extraction and marker genotyping (*43*).

### Synergy of long-standing germplasm exchange practices with new genomics technologies

In this study, we integrated evolutionary population genomic analyses and historical records on global germplasm exchange to show that the recent evolutionary rescue of sorghum depended on >50 years of germplasm exchange. Germplasm exchange led to global diffusion of a rare SCA resistance allele, sometimes purposely and sometimes inadvertently, from smallholder farmers in Ethiopian highlands across breeding programs in Africa, the Americas, and Asia. Over the past several decades, movement of crop genetic resources through international cooperation of germplasm exchange has provided access to adaptive genetic variation for crop improvement (*31*). However, germplasm exchange is increasingly restricted due to commercial or institutional interests asserting intellectual property (IP) rights and governments asserting national or local sovereignty over genetic resources (*44*). While IP rights and sovereignty are important considerations, the question remains how to balance these aims with the benefits of free exchange of global public goods (*45, 46*). While we are not in a position to resolve these societal tradeoffs, our study does highlight the global food security benefits of germplasm exchange and the opportunities that could be lost due to restrictions on exchange.

Taken together, our findings suggest that new genomic technologies will be most powerful when leveraged with global exchange of germplasm and knowledge. No matter how powerful new genomic technologies are in terms of accuracy or throughput, their utility will depend on the germplasm assayed, since all genetic mapping approaches require effective recombination and allelic diversity (*47, 48*). Global germplasm exchange vastly increases both these parameters, providing a “bank” of historical recombinations and allelic variants that can be rapidly leveraged with new genomic tools (Fig. 3, 4). Therefore, our best opportunity to address challenges of global change may be to leverage new genomic technologies with long-standing practices of global germplasm exchange.

## MATERIALS AND METHODS

### Sorghum breeding and production in Haiti

The Chibas sorghum breeding program was launched in 2013 using admixed global germplasm, including heterogeneous breeding material from West Africa carrying *ms3* nuclear male sterility, and inbred global accessions. During 2015–2018, the material was selected in breeding nurseries under low-input conditions (approximating local smallholder practices) and extensively intercrossed using the *ms3* sterility system. No insecticides were used to limit SCA infestations in breeding nurseries in this period and natural SCA infestations were intense during this period (e.g. Fig. 1C). Note, selection pressure on sorghum by SCA in Haiti is expected to be greater than in temperate zone (e.g. U.S.) because the SCA infestation occurs year-round in this tropical environment. Annual sorghum production estimates for Haiti are based on FAOSTAT (2009– 2014 and 2018) (*49*) and the USDA forecast for 2019-2020 (*19*). FAOSTAT data for 2015–2017 and 2019 was not used because it was based on imputation (“FAO data based on imputation methodology”) that did not account for the known effects of SCA (e.g. “this aphid spread throughout the country and decimated Haiti sorghum production”) (*19*). Production for the missing years of SCA outbreak was inferred based on 2009 agriculture survey acreage prior to infestation in each region and assessment of sorghum production levels compared to pre-infestation levels, adjusted to FAOSTAT (1990–2014) production averages for each region.

### Plant genetic resources

The HBP (N = 296) are inbred lines derived from a recurrent selection breeding population developed by intercrossing germplasm that survived natural SCA infestation. For genomic DNA extraction, fresh leaf tissue of each accession was collected from two weeks old seedlings raised in a greenhouse. Tissue was lyophilized for two days and then grounded up using a 96-well plate plant tissue grinder (Retsch Mixer Mill). Genomic DNA was extracted using the BioSprint 96 DNA Plant Kit (QIAGEN), quantified using Quant-iT™ PicoGreen® dsDNA Assay Kit, and normalized to 10 ng/uL. An additional set of global accessions (GDP, *N* = 767) was assembled based on a published data set (*50, 51*) including sorghum accessions from 52 countries on five continents and all major botanical types (Supp. Fig. S1, Supp. File S1). The GDP accessions included 164 caudatum, 96 guinea, 81 durra, 57 bicolor, and 47 kafir accessions, along with 288 of other botanical types and 34 accessions of unknown botanical type.

### Genotyping-by-sequencing

Genotypes for the 296 Haitian breeding lines were generated with genotyping-by-sequencing. Genomic DNA digestion, ligation and PCR amplification processes were performed according to the methods previously described (*50*). The libraries were sequenced using the single-end 100-cycle sequencing by Illumina HiSeq2500 (Illumina, San Diego CA, USA) at the University of Kansas Medical Center, Kansas City, MO, USA. A total of 220 million reads for the HBP were combined with published data for the GDP (*50*) for SNP calling. TASSEL 5 GBS v2 pipeline (*52*) was used to perform the SNP calling of the sequence data obtained from Illumina sequencing. Reads were aligned to the BTx623 sorghum reference genome v.3.1 (*53*) with the Burrows-Wheeler Alignment (*54*). The SNPs were filtered for 20% missingness, then missing data were imputed using BEAGLE 4.0 (*55*). Genotyping data are available at Dryad [*accession to be added following acceptance*].

### Population genomic analyses

Genome-wide nucleotide diversity (π) and Tajima’s *D* statistics for HBP and GDP were estimated based on a non-overlapping sliding window of 1 Mbp across the genome using VCFtools (*56*). The characterization of the population structure of the HBP was based on a discriminant analysis of principal components (DAPC) in the Adegenet package in R (*57*). A distance matrix calculated based on a modified Euclidean distance model was used to create a cladogram based on a neighbor-joining algorithm in TASSEL (*58*). Neighbor-joining analysis was visualized using the APE package in R (*59*). The population structure of the germplasm panel was further assessed by the Bayesian model-based clustering method implemented in the ADMIXTURE program (*60*). Pairwise SNP differentiation (*F*_ST_) between the HBP and the GDP were calculated and outlier loci were detected based on an inferred distribution of neutral *F*_ST_ using the R Package OutFLANK (*61*).

### Whole genome resequencing

Around the 130 kb mapped interval in BTx623, SNPs from 10 sorghum accessions with known SCA resistance status were examined to search for functional mutations responsible for SCA resistance. Six of the 10 resequenced accessions represent known susceptible lines, which include RTx430 (PI 655996), BTx623 (PI 564163), Tx7000 (PI 655986), Tx2737 (PI 655978), BTx642, and RTx436. The remaining four resequenced accessions represent known resistant lines, which includes PI 257599 (SC110 original exotic parent), PI 276837 (SC170 original exotic parent), PI 534157 (SC170), and IS 36563 (IRAT204). These samples were used pre-publication for this interval analysis with permission from TERRA-REF (Mockler), JGI Sorghum Pan-genome project (Mockler), BMFG Sorghum Genomic Toolbox (Mockler and Morris), JGI Sorghum Diversity project (John Mullet), and the JGI EPICON project (Vogel). The reads were mapped to *Sorghum bicolor* v3.1 using bwa-mem. The bam file was filtered for duplicates using Picard (http://broadinstitute.github.io/picard) and realigned around indels using GATK (*62*). Multi-sample SNP calling was done using SAMtools mpileup and Varscan V2.4.0 with a minimum coverage of 8 and a minimum alternate allele frequency of four. Repeat content of the genome was masked using 24 bp kmers. Kmers that occur at a high frequency, up to 5%, were masked. SNPs around 25 bp of the mask were removed for further analysis. A SNP was included for further analysis only when it has coverage in 75% of the samples, and a MAF > 0.005. Functional annotation of the variants within the *RMES1* locus was performed using SNPEff.

### *De novo* genome sequencing

*De novo* genome assembly of the resistance sorghum line PI 276837 was used to perform comparative genomic analysis to identify the causative variant for SCA resistance at the *RMES1* locus. PI 276837 main assembly consisted of 101.47x of PACBIO coverage with an average read size of 11,931 bp. The genome was assembled using Canu 1.8, a fork of the Celera Assembler designed for high-noise single-molecule sequencing. The resulting sequence was polished using ARROW. The assembled genome resulted in contig N50 sizes ranging from 14 to 19 kb and scaffold N50 sizes ranging from 5 to 65 kb. Sequence variations at *RMES1* locus between the *de novo* sequence of PI 276837 were compared to the reference genomes of BTx623, Tx430, and BTx642.

### KASP marker development

SNPs from the *F*_ST_ genomic selection scan were selected for development into markers based on several factors: LOD score of the *F*_ST_ analysis, proximity to *RMES1* locus, and suitability of the flanking sequence for KASP assay development. The KASP assays were developed utilizing a third-party genotyping service provider, Intertek AgriTech (Alnarp, Sweden), who designed the KASP assays via the Kraken software. All genomic DNA extraction and KASP genotyping were performed by Intertek using two 6 mm leaf punches dried with silica beads. Initial technical validation of the KASP marker was performed using known resistant (SC110, IRAT204 and Tx2783) and susceptible (KS585 and Tx7000) sorghum lines. Further validation was performed by genotyping a panel of 10 known resistant and 28 known susceptible lines, along with multiple F1 crosses of each of the lines. The KASP markers developed for SCA resistance selection (Supp. File S6) are publicly available through the third-party genotyping service provided by Intertek. For further information on accessing markers contact the corresponding author.

### Marker validation in public and commercial breeding programs

To test the predictiveness of the marker, a population segregating for SCA resistance was developed by crossing the susceptible Tx430 and resistant IRAT204. F_3_ and F_4_ lines of the Tx430 x IRAT204 population were genotyped with the KASP marker together with the susceptible and resistant parents. The same population was evaluated for SCA reaction using a free-choice flat-screen trial in the greenhouse. Tx2783 and SC110 were included as known resistant genotypes, along with the known susceptible genotypes, KS 585 and Tx7000 (*63, 64*). Free-choice flat-screen assay, data collection (damage rating, SPAD score, and plant height difference), and analysis were conducted as previously described (*24*). Validation of the KASP marker across different breeding programs was performed in eight breeding programs, five commercial and three public in the US. Each program collected tissue samples from known tolerant and susceptible parental breeding lines, F_1_s of the parental lines, and later generation lines from their SCA tolerance breeding populations; the SCA reaction phenotypes of the late generation lines may or may not have been known. For the parental breeding lines, both technical replicates (tissue samples from the same plant) and biological replicates (tissue samples from separate plants) were collected in order to test both the technical function of the markers and the reliability of the germplasm, respectively. Additionally, most programs included public sources (e.g. Tx2783) of known SCA tolerance as checks. Tissue samples were sent to Intertek, who extracted DNA and performed the KASP genotyping.

## ACKNOWLEDGMENTS

This study is made possible by the support of the American People provided to the Feed the Future Innovation Lab for Collaborative Research on Sorghum and Millet through the United States Agency for International Development (USAID) under Associate Award No. AID-OAA-LA-16-00003, “Feed the Future Innovation Lab for Genomics-Assisted Sorghum Breeding”. The contents are the sole responsibility of the authors and do not necessarily reflect the views of USAID or the United States Government. The work conducted by the US Department of Energy Joint Genome Institute is supported by the Office of Science of the US Department of Energy under Contract No DE-AC02-05CH11231. We thank the Joint Genome Institute and collaborators for pre-publication access to the genomes of Tx430 and Tx642 for use in this study. Additional support was provided by the Bill & Melinda Gates Foundation under the “Sorghum Genomics Toolkit” project. We thank the breeding programs that participated in the marker testing. We thank Matt Davis for technical support and Jesse Lasky for comments on the manuscript. Such use does not constitute an official endorsement or approval by the United States Department of Agriculture or the Agricultural Research Service of any product or service to the exclusion of others that may be suitable. USDA is an equal opportunity provider and employer. The authors declare that they have no competing interests. All data needed to evaluate the conclusions in the paper are present in the Supplementary Materials or Dryad.

