## Supplementary material for "The recent evolutionary rescue of a staple crop depended on over half a century of global germplasm exchange": Supp. Fig.

### Slide 1
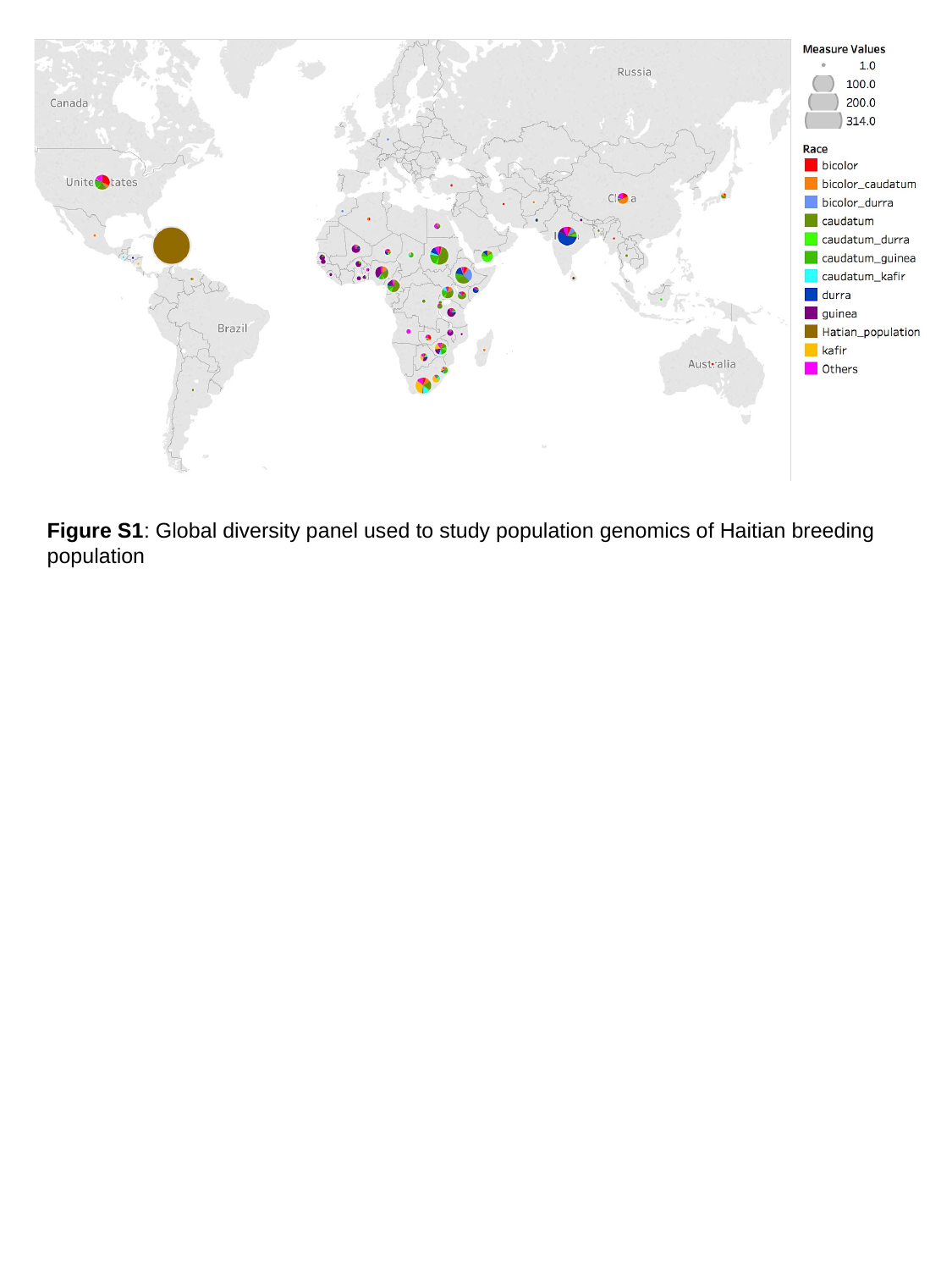

Figure S1: Global diversity panel used to study population genomics of Haitian breeding population

### Slide 2
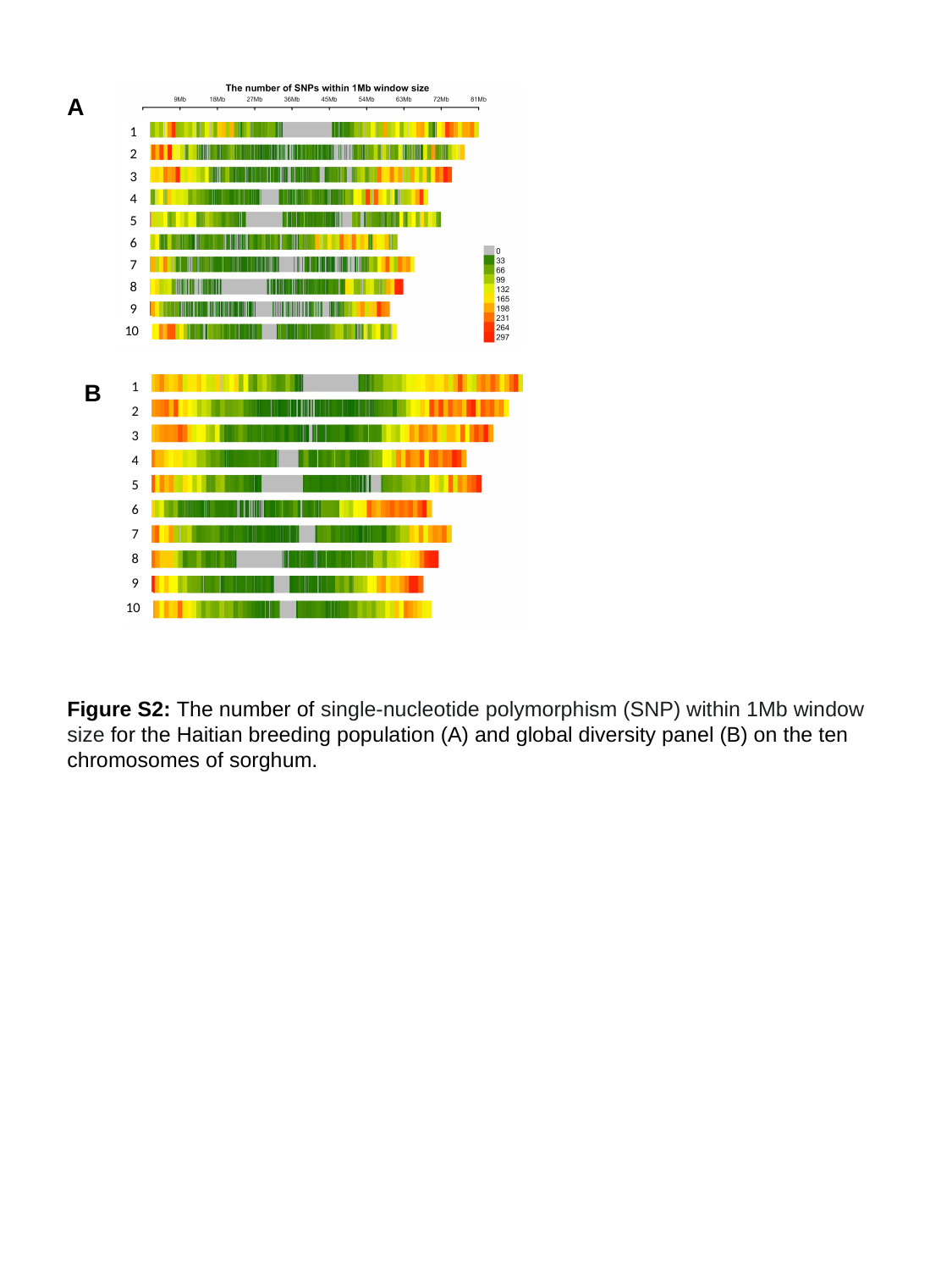

1
2
3
4
5
6
7
8
9
10
A
1
2
3
4
5
6
7
8
9
10
B
Figure S2: The number of single‐nucleotide polymorphism (SNP) within 1Mb window size for the Haitian breeding population (A) and global diversity panel (B) on the ten chromosomes of sorghum.

### Slide 3
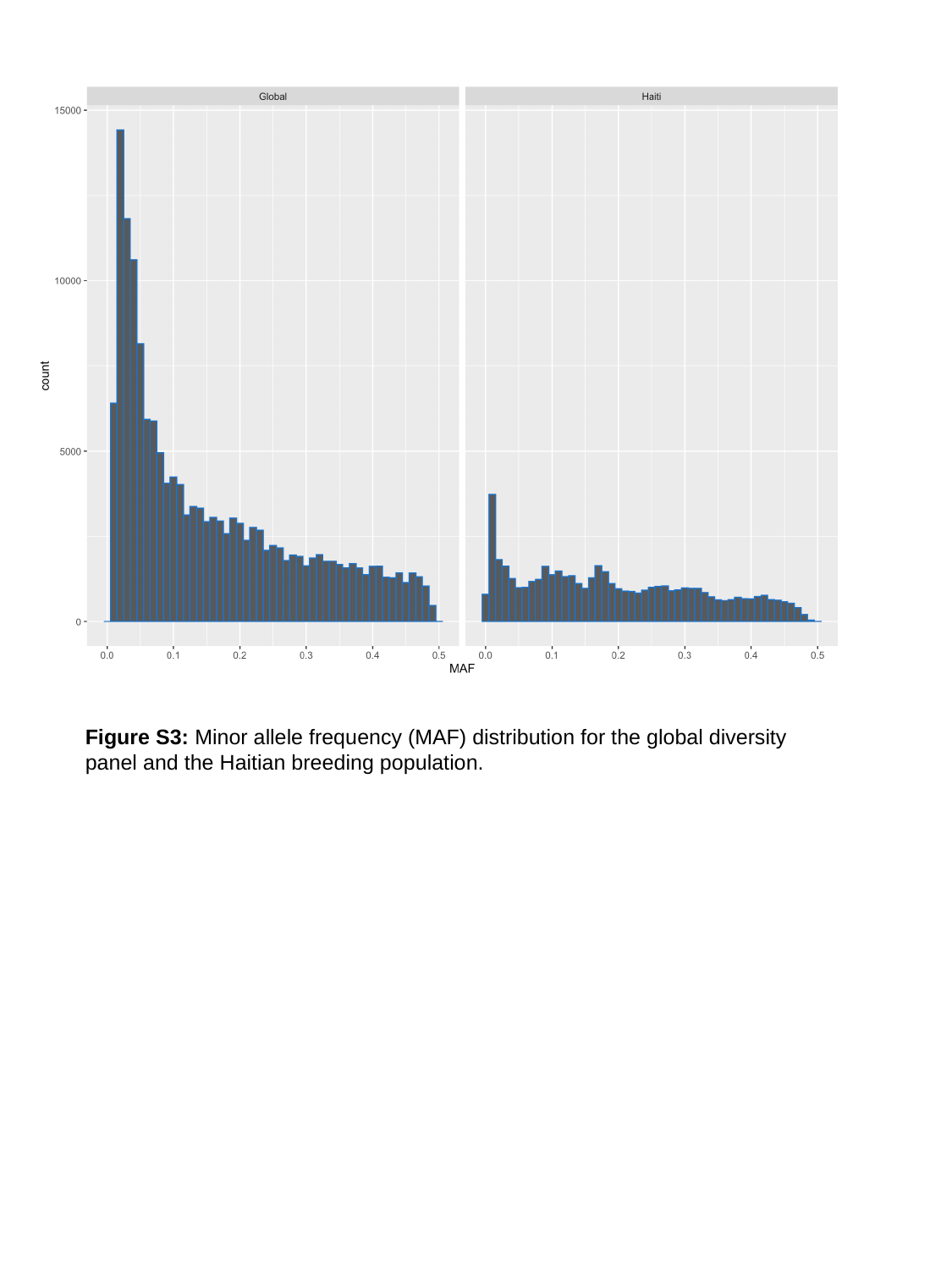

Figure S3: Minor allele frequency (MAF) distribution for the global diversity panel and the Haitian breeding population.

### Slide 4
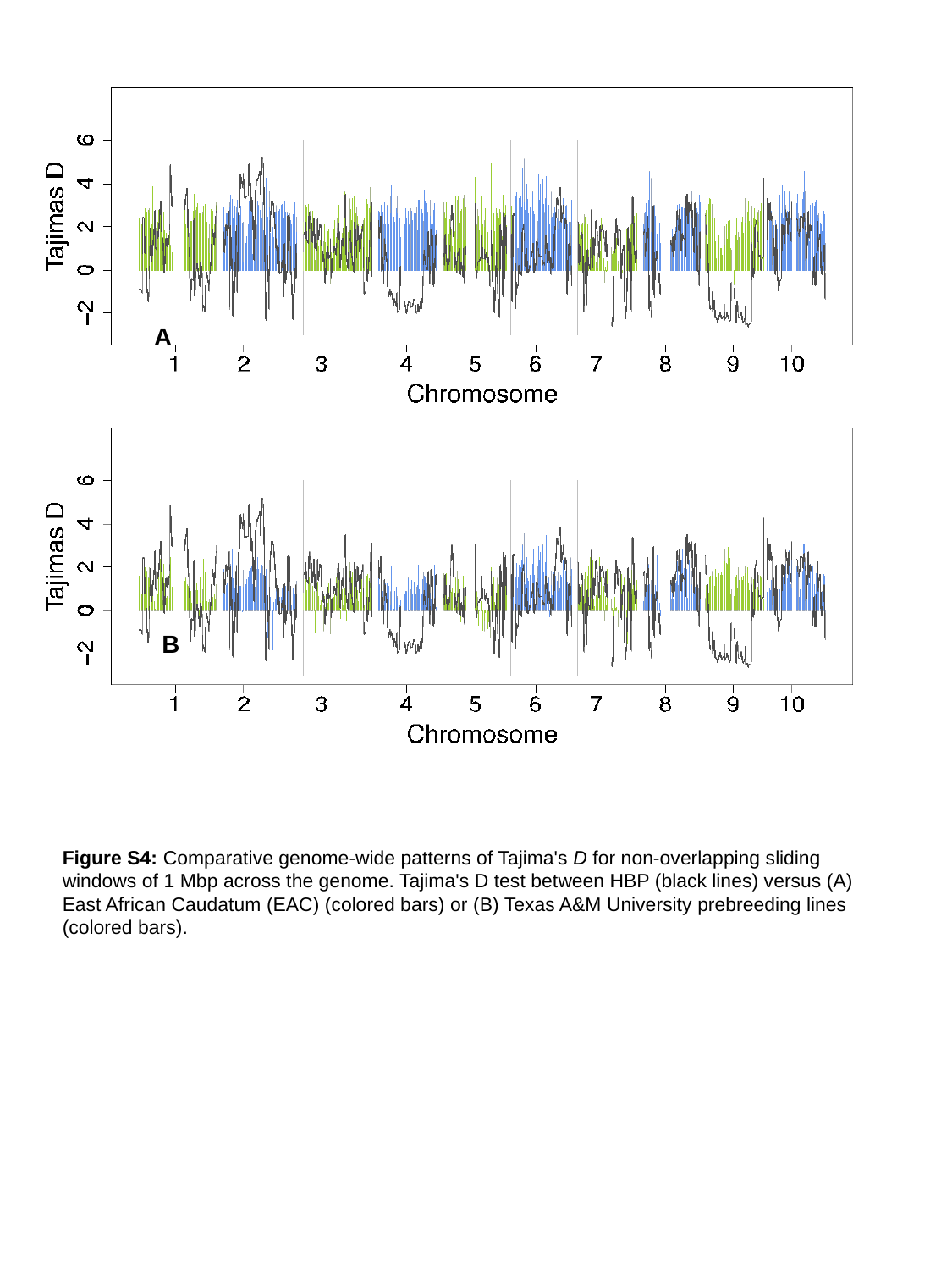

A
B
Figure S4: Comparative genome-wide patterns of Tajima's D for non-overlapping sliding windows of 1 Mbp across the genome. Tajima's D test between HBP (black lines) versus (A) East African Caudatum (EAC) (colored bars) or (B) Texas A&M University prebreeding lines (colored bars).

### Slide 5
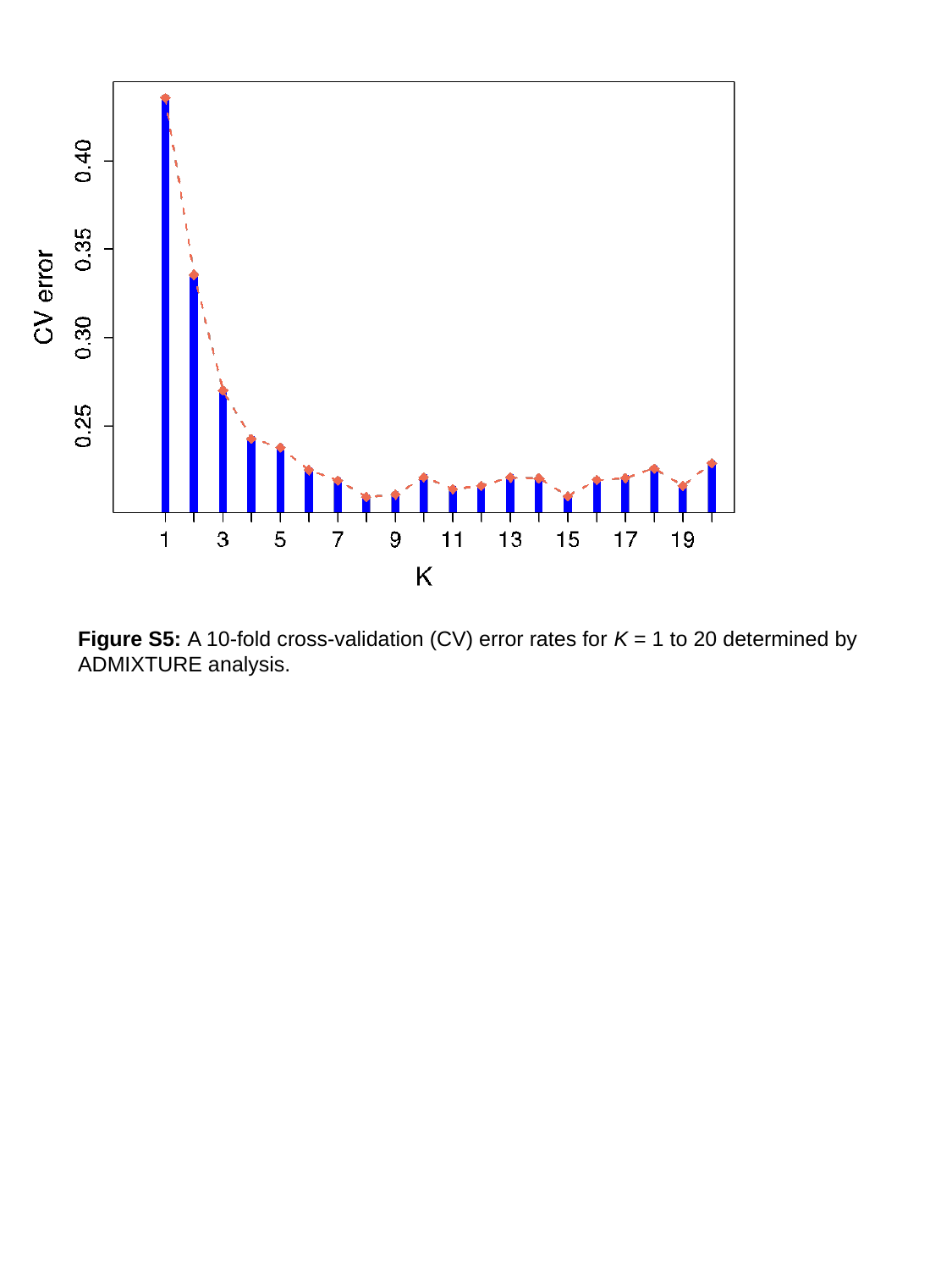

Figure S5: A 10-fold cross-validation (CV) error rates for K = 1 to 20 determined by ADMIXTURE analysis.

### Slide 6
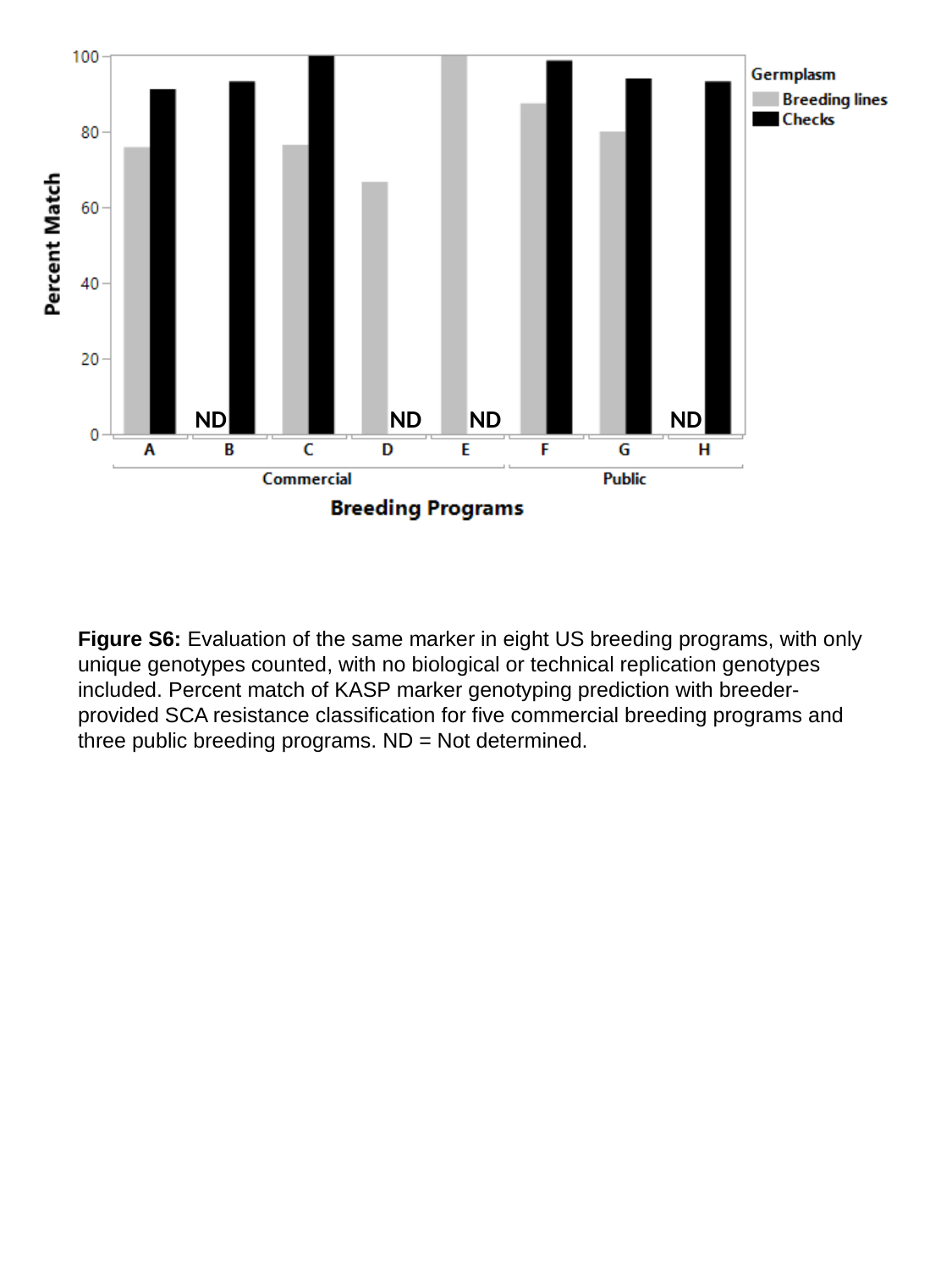

ND
ND
ND
ND
Figure S6: Evaluation of the same marker in eight US breeding programs, with only unique genotypes counted, with no biological or technical replication genotypes included. Percent match of KASP marker genotyping prediction with breeder-provided SCA resistance classification for five commercial breeding programs and three public breeding programs. ND = Not determined.

### Slide 7
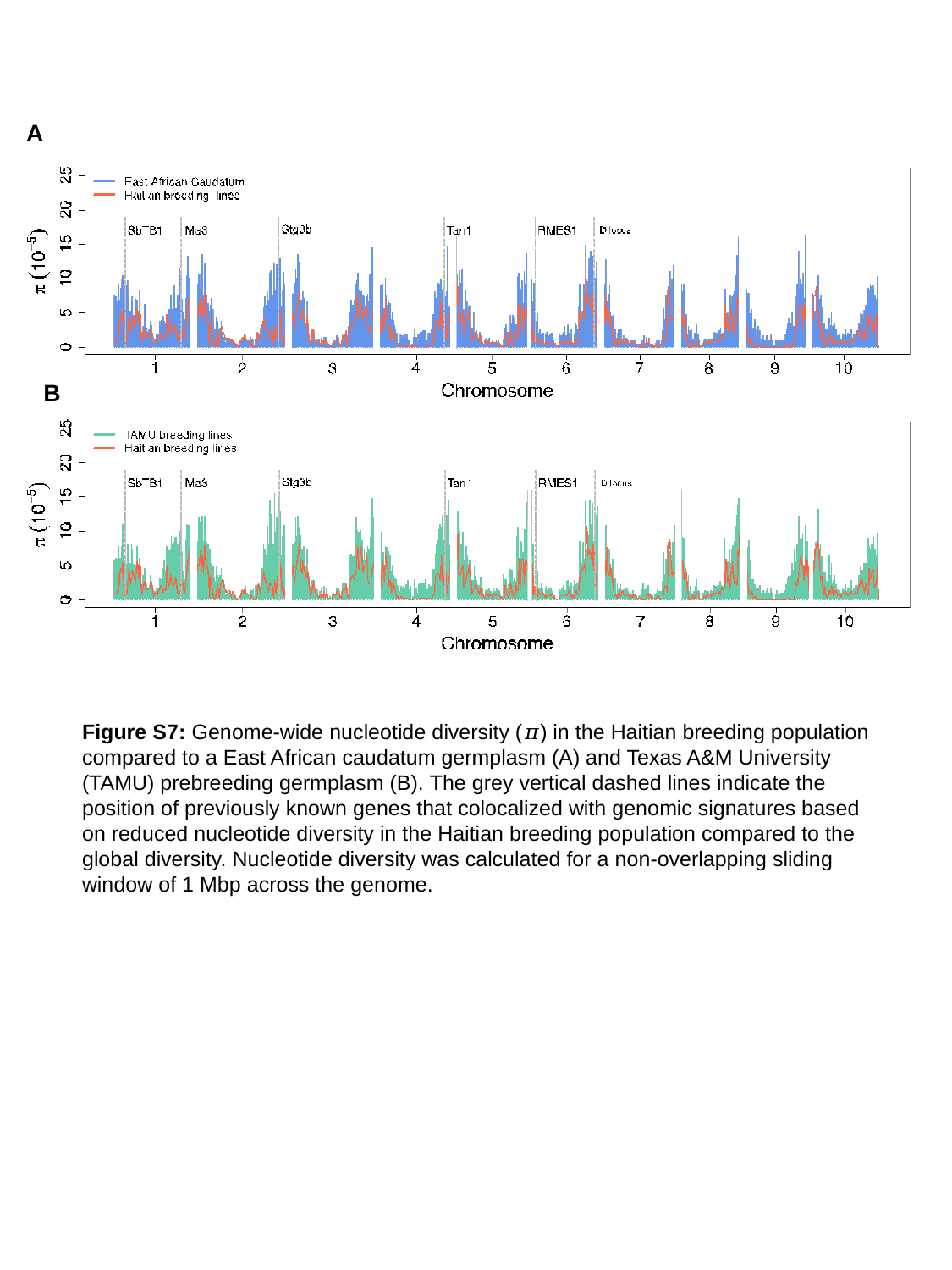

A
B
Figure S7: Genome-wide nucleotide diversity (𝜋) in the Haitian breeding population compared to a East African caudatum germplasm (A) and Texas A&M University (TAMU) prebreeding germplasm (B). The grey vertical dashed lines indicate the position of previously known genes that colocalized with genomic signatures based on reduced nucleotide diversity in the Haitian breeding population compared to the global diversity. Nucleotide diversity was calculated for a non-overlapping sliding window of 1 Mbp across the genome.

### Slide 8
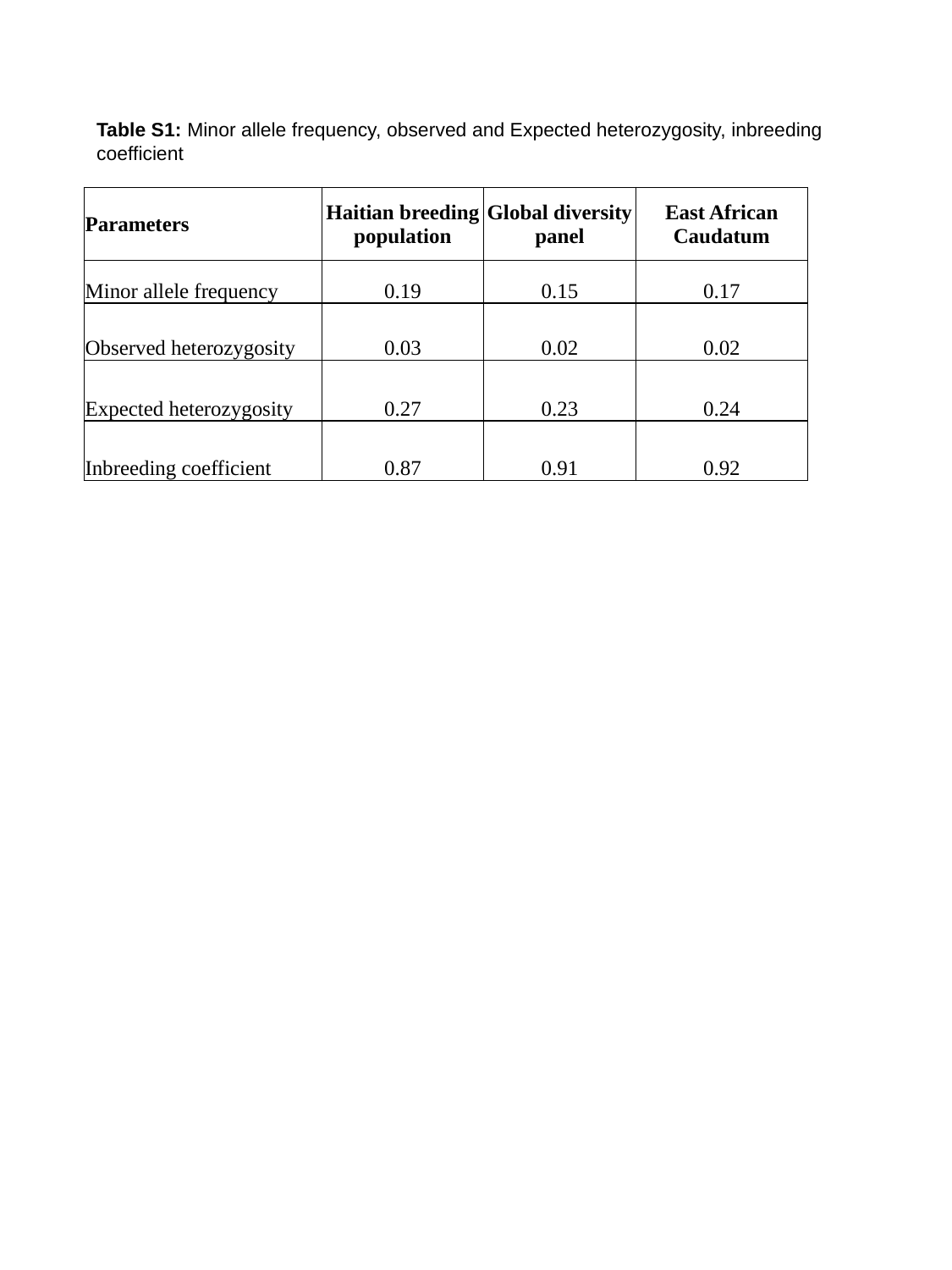

Table S1: Minor allele frequency, observed and Expected heterozygosity, inbreeding coefficient
| Parameters | Haitian breeding population | Global diversity panel | East African Caudatum |
| --- | --- | --- | --- |
| Minor allele frequency | 0.19 | 0.15 | 0.17 |
| Observed heterozygosity | 0.03 | 0.02 | 0.02 |
| Expected heterozygosity | 0.27 | 0.23 | 0.24 |
| Inbreeding coefficient | 0.87 | 0.91 | 0.92 |

### Slide 9
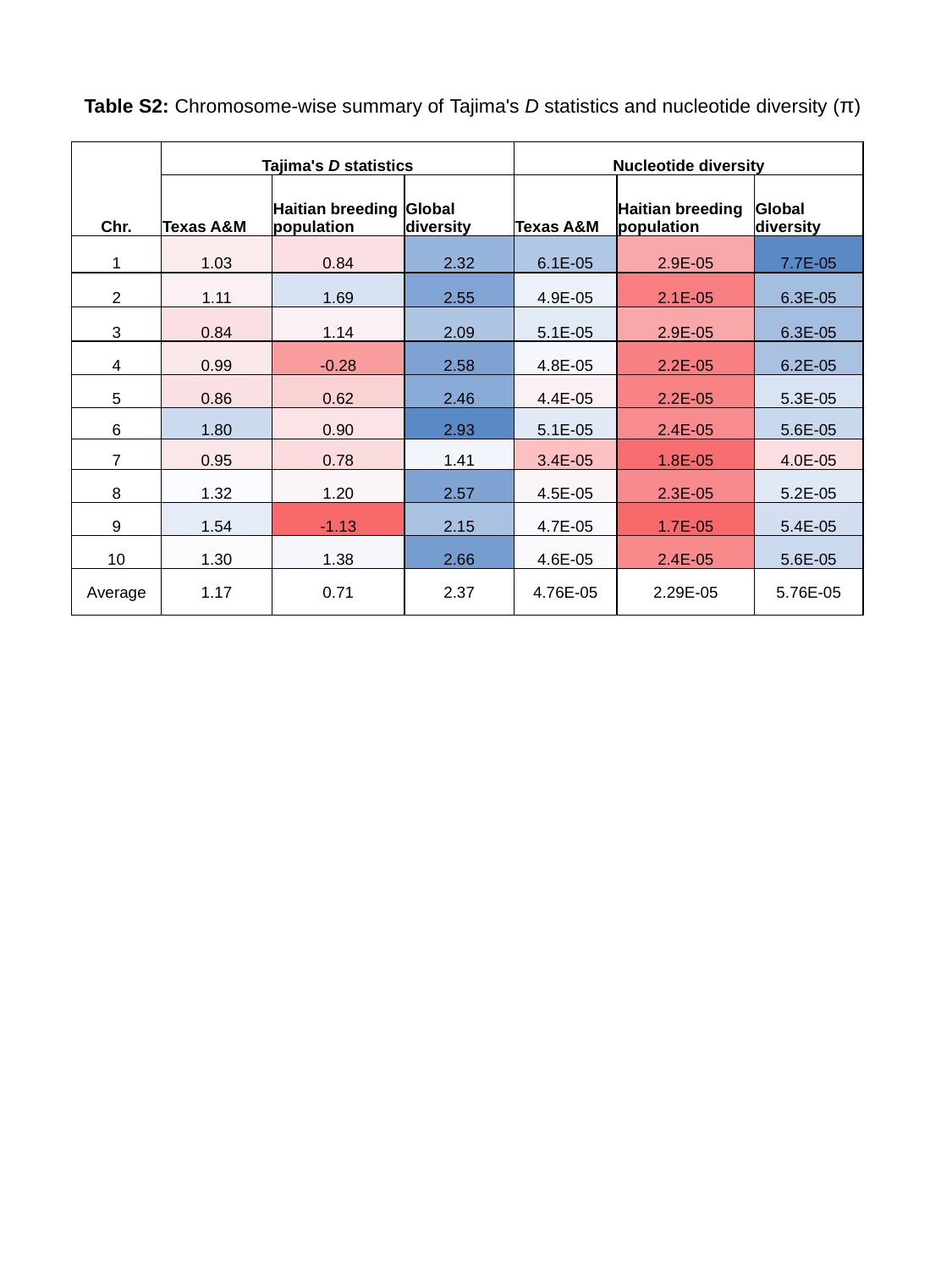

Table S2: Chromosome-wise summary of Tajima's D statistics and nucleotide diversity (π)
| Chr. | Tajima's D statistics | | | Nucleotide diversity | | |
| --- | --- | --- | --- | --- | --- | --- |
| | Texas A&M | Haitian breeding population | Global diversity | Texas A&M | Haitian breeding population | Global diversity |
| 1 | 1.03 | 0.84 | 2.32 | 6.1E-05 | 2.9E-05 | 7.7E-05 |
| 2 | 1.11 | 1.69 | 2.55 | 4.9E-05 | 2.1E-05 | 6.3E-05 |
| 3 | 0.84 | 1.14 | 2.09 | 5.1E-05 | 2.9E-05 | 6.3E-05 |
| 4 | 0.99 | -0.28 | 2.58 | 4.8E-05 | 2.2E-05 | 6.2E-05 |
| 5 | 0.86 | 0.62 | 2.46 | 4.4E-05 | 2.2E-05 | 5.3E-05 |
| 6 | 1.80 | 0.90 | 2.93 | 5.1E-05 | 2.4E-05 | 5.6E-05 |
| 7 | 0.95 | 0.78 | 1.41 | 3.4E-05 | 1.8E-05 | 4.0E-05 |
| 8 | 1.32 | 1.20 | 2.57 | 4.5E-05 | 2.3E-05 | 5.2E-05 |
| 9 | 1.54 | -1.13 | 2.15 | 4.7E-05 | 1.7E-05 | 5.4E-05 |
| 10 | 1.30 | 1.38 | 2.66 | 4.6E-05 | 2.4E-05 | 5.6E-05 |
| Average | 1.17 | 0.71 | 2.37 | 4.76E-05 | 2.29E-05 | 5.76E-05 |

### Slide 10
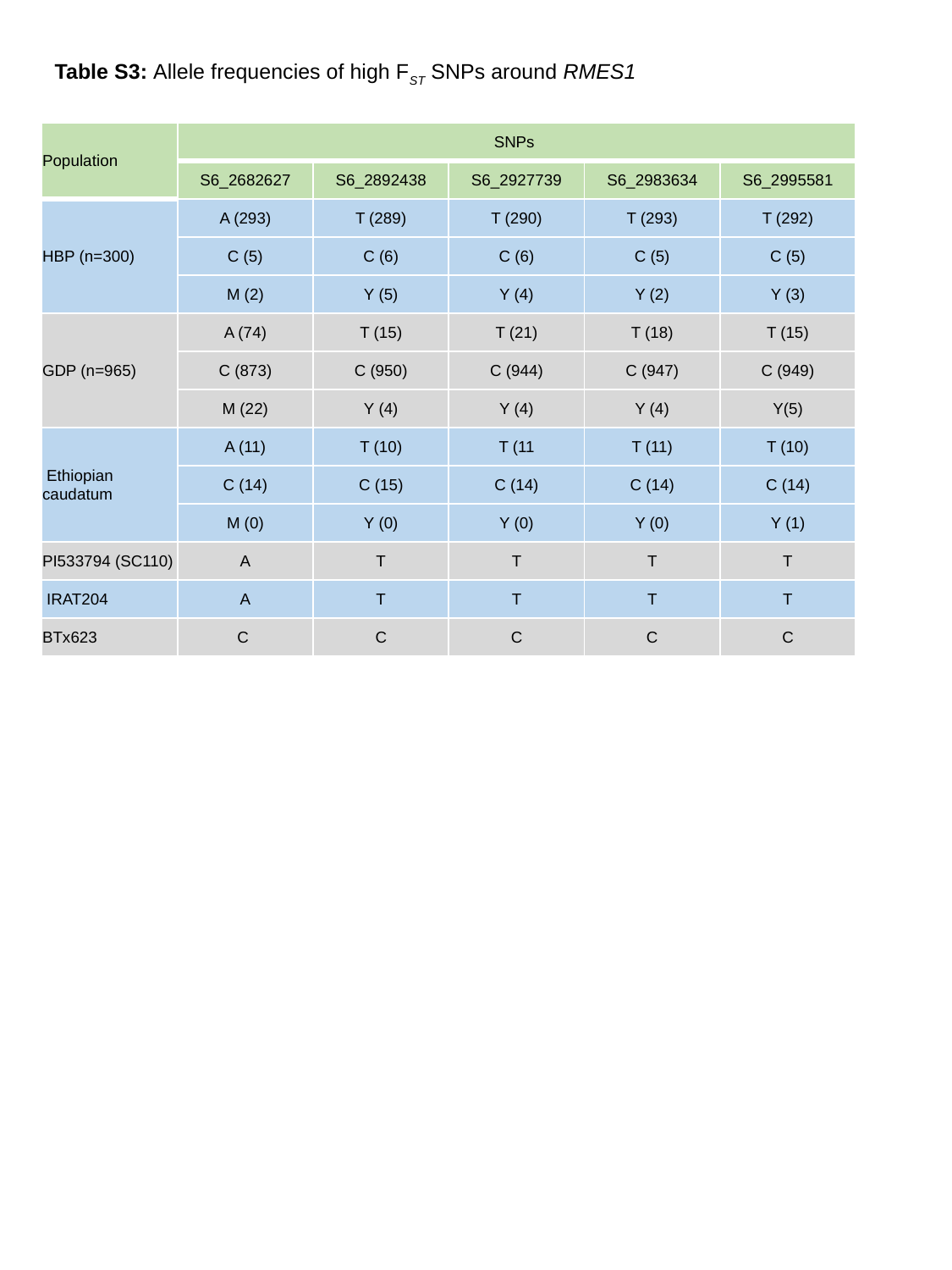

Table S3: Allele frequencies of high F_ST_ SNPs around RMES1
| Population | SNPs | | | | |
| --- | --- | --- | --- | --- | --- |
| | S6\_2682627 | S6\_2892438 | S6\_2927739 | S6\_2983634 | S6\_2995581 |
| HBP (n=300) | A (293) | T (289) | T (290) | T (293) | T (292) |
| | C (5) | C (6) | C (6) | C (5) | C (5) |
| | M (2) | Y (5) | Y (4) | Y (2) | Y (3) |
| GDP (n=965) | A (74) | T (15) | T (21) | T (18) | T (15) |
| | C (873) | C (950) | C (944) | C (947) | C (949) |
| | M (22) | Y (4) | Y (4) | Y (4) | Y(5) |
| Ethiopian caudatum | A (11) | T (10) | T (11 | T (11) | T (10) |
| | C (14) | C (15) | C (14) | C (14) | C (14) |
| | M (0) | Y (0) | Y (0) | Y (0) | Y (1) |
| PI533794 (SC110) | A | T | T | T | T |
| IRAT204 | A | T | T | T | T |
| BTx623 | C | C | C | C | C |

### Slide 11
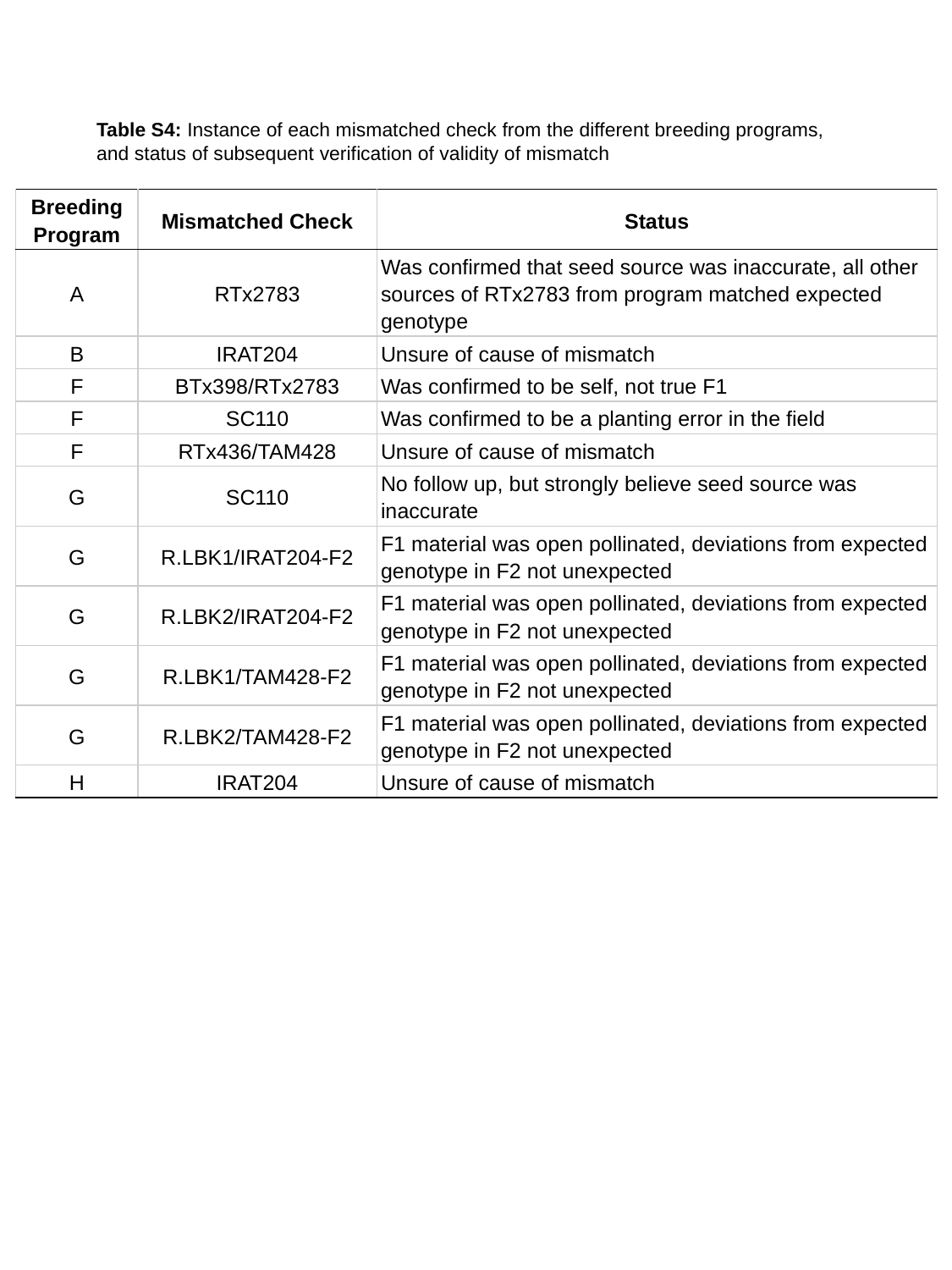

Table S4: Instance of each mismatched check from the different breeding programs, and status of subsequent verification of validity of mismatch
| Breeding Program | Mismatched Check | Status |
| --- | --- | --- |
| A | RTx2783 | Was confirmed that seed source was inaccurate, all other sources of RTx2783 from program matched expected genotype |
| B | IRAT204 | Unsure of cause of mismatch |
| F | BTx398/RTx2783 | Was confirmed to be self, not true F1 |
| F | SC110 | Was confirmed to be a planting error in the field |
| F | RTx436/TAM428 | Unsure of cause of mismatch |
| G | SC110 | No follow up, but strongly believe seed source was inaccurate |
| G | R.LBK1/IRAT204-F2 | F1 material was open pollinated, deviations from expected genotype in F2 not unexpected |
| G | R.LBK2/IRAT204-F2 | F1 material was open pollinated, deviations from expected genotype in F2 not unexpected |
| G | R.LBK1/TAM428-F2 | F1 material was open pollinated, deviations from expected genotype in F2 not unexpected |
| G | R.LBK2/TAM428-F2 | F1 material was open pollinated, deviations from expected genotype in F2 not unexpected |
| H | IRAT204 | Unsure of cause of mismatch |
